# Reference-based comparison of adaptive immune receptor repertoires

**DOI:** 10.1101/2022.01.23.476436

**Authors:** Cédric R. Weber, Teresa Rubio, Longlong Wang, Wei Zhang, Philippe A. Robert, Rahmad Akbar, Igor Snapkov, Jinghua Wu, Marieke L. Kuijjer, Sonia Tarazona, Ana Conesa, Geir K. Sandve, Xiao Liu, Sai T. Reddy, Victor Greiff

## Abstract

B- and T-cell receptor (immune) repertoires can represent an individual’s immune history. While current repertoire analysis methods aim to discriminate between health and disease states, they are typically based on only a limited number of parameters (e.g., clonal diversity, germline usage). Here, we introduce immuneREF: a quantitative multi-dimensional measure of adaptive immune repertoire (and transcriptome) similarity that allows interpretation of immune repertoire variation by relying on both repertoire features and cross-referencing of simulated and experimental datasets. immuneREF is implemented in an R package and was validated based on detection sensitivity of immune repertoires with known similarities and dissimilarities. To quantify immune repertoire similarity landscapes across health and disease, we applied immuneREF to >2400 datasets from individuals with varying immune states (healthy, [autoimmune] disease and infection [Covid-19], immune cell population). Importantly we discovered, in contrast to the current paradigm, that blood-derived immune repertoires of healthy and diseased individuals are highly similar for certain immune states, suggesting that repertoire changes to immune perturbations are less pronounced than previously thought. In conclusion, immuneREF implements population-wide analysis of immune repertoire similarity and thus enables the study of the adaptive immune response across health and disease states.

## Introduction

B- and T-cell receptor (BCR, TCR) repertoires (also called adaptive immune receptor repertoires, AIRR) are continually shaped throughout the lifetime of an individual in response to environmental and pathogenic exposure. As of yet, however, there exists only a limited quantitative conception of how immune receptor repertoires differ across individuals and cell populations (Brown et al., 2019; Miho et al., 2018; Raybould et al., 2021). This is primarily because a method for measuring inter-individual (inter-repertoire) similarity is lacking, thus greatly impeding the understanding of how health and disease shape immune repertoires and how disease contributes to the deviation of an individuals’ baseline repertoire (Cobey et al., 2015). Although it is generally thought that infection or disease induce measurable repertoire changes (even on the antigen-specific agnostic level), this belief remains unproven and, in fact, is counter to current evidence finding, using statistical learning, that even in systemic infections such as CMV only a comparatively very small number of TCRs are infection associated (DeWitt et al., 2018; Emerson et al., 2017; Pavlović et al., 2021). As opposed to machine learning approaches that aim to detect the most differentiating factors (i.e., subsets of a repertoire) between, for example, two different immune states (Greiff et al., 2020; Pavlović et al., 2021; Pertseva et al., 2021; Shemesh et al., 2021; Widrich et al., 2020a, 2020b), we investigate here a method for quantitatively comparing any two repertoires in an unsupervised fashion. We thus seek to understand to what extent individuals differ with respect to their entire repertoire and not just class-associated subsets.

The need for comparing immune repertoires using a quantitative measure has recently been addressed by approaches based on single sequence-dependent and sequence-independent features, which vary in statistical dependency (mutual information) and immunological interpretability (Chiffelle et al., 2020; Miho et al., 2018; Olson et al., 2019). Sequence-dependent approaches range from the measurement of clonal overlap (Bolen et al., 2017; Greiff et al., 2015a; Miho et al., 2018; Yaari and Kleinstein, 2015) to more sophisticated algorithms that identify disease-specific enrichment of sequence clusters by testing against VDJ recombination models (Pogorelyy et al., 2019) or similarity networks of control datasets (Pogorelyy and Shugay, 2019; Shugay et al., 2015). Sequence-independent approaches are mainly represented by entropy-based diversity indices (Alon et al., 2021; Greiff et al., 2015a; Kaplinsky and Arnaout, 2016; Strauli and Hernandez, 2016), which have lately been augmented with a correction for sequence similarity (Arora et al., 2018; Vujović et al., 2021). None of the currently available comparative methods, which are based on single repertoire features, however, represent an integrated multi-feature measure of immune repertoire similarity that takes into account the complexity of information encoded in the ensemble of the existing immune repertoire features (Gupta et al., 2015; Heiden et al., 2014; Nazarov et al., 2020; Shugay et al., 2015). Such an integrated measure, encoding per-feature similarity in one common mathematical structure, is needed to enable a representation of repertoire similarity.

Here, we introduce immuneREF: a measure for quantifying immune repertoire similarity across multiple immune repertoire features. Our framework, implemented in an R package, measures immune repertoire similarity using a combination of features that are immunologically interpretable (clonal expansion, sequence composition, repertoire architecture, and clonal overlap) and that cover largely distinct dimensions of the immune repertoire spaces. Specifically, to interpret immune repertoire similarity scores, immuneREF establishes a self-augmenting dictionary of simulated and experimental datasets where each new dataset analyzed may be used as a comparative reference for scoring and biologically interpreting inter-individual variation (and thus the deviation) of immune repertoire features (Fig. 1, Supplementary Fig. 1). We applied immuneREF to >2400 immune repertoires from humans with varying immune states (healthy, virus infection, autoimmune disease) and found that the similarity of blood-derived immune repertoires is not consistently a function of the immune state.

**Figure 1.**
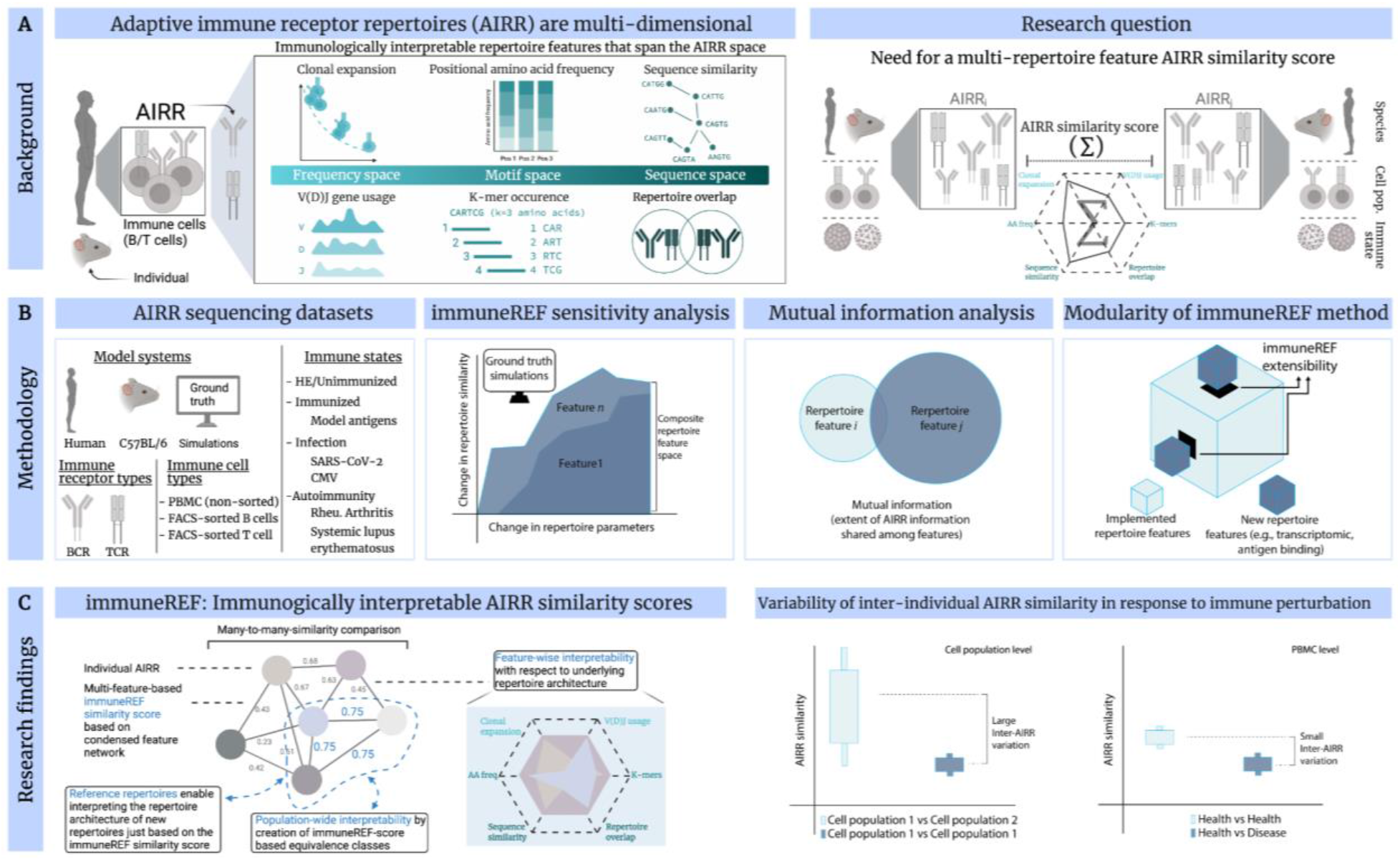
Reference-based comparison of adaptive immune receptor repertoires (AIRRs). **(A)** The complexity of AIRRs spans the frequency, motif, and feature space to each of which distinct repertoire features may be attributed: the immune information stored in AIRRs is multidimensional. A longstanding question in the AIRR field is how to quantitatively measure inter-sample (sample: e.g., individual, immune cell population) AIRR similarity by accounting for AIRR feature multidimensionality in the effort to understand the distribution of inter-sample AIRR similarity across different immune events or immune cell populations. **(B)** We set out to develop an AIRR similarity measure that is sensitive, captures maximal immune information, and is sufficiently flexible to allow future integration of additional repertoire features (extensibility). **(C)** Each AIRR is represented as a node in a similarity network. The edges connecting the nodes represent the similarity score between the AIRR based on the six repertoire features. The immuneREF approach establishes interpretability on different levels: (i) from a single-feature perspective, the application of spider plots allows for an interpretable comparative analysis between repertoires, enabling the user to interpret the result observed in the condensed network on a per feature basis. (ii) From the condensed feature network perspective, a major novelty introduced by the immuneREF workflow is the ability to combine established repertoire features into a common coordinate system. This transformation allows the combination of trends across features into a single condensed network that represents pairwise-cross-feature similarities. These pairwise similarities allow for the identification of subsets of more similar or aberrant repertoires. Interpretability on both features means allowing comparison to other repertoires, and to simulated ones (of which we know the repertoire structure as ground truth) thus creating similarity equivalence classes. Equivalence classes create sets of reference repertoires, which enable interpreting the repertoire structures of other repertoires solely based on the immuneREF similarity score.

Overall, immuneREF enables the quantification of repertoire similarity at population scale while still providing single-individual resolution and enables answering of fundamental questions such as to what extent immune repertoires are robust to perturbations introduced by immune events.

## Methods

### immuneREF features

For each dataset, we calculated six immune repertoire features and a per-feature similarity score.

#### immuneREF Feature: Evenness profiles (state of clonal expansion)

Evenness profiles were calculated as described previously (Greiff et al., 2015b) on the CDR3 nucleotide level. Briefly, we calculated the Hill-diversity for alpha values 0–10 in steps of 0.1 with alpha = 1 being defined as the Shannon evenness. Each entry in the profile varies between ≈0 and 1, where higher values indicate an increasingly uniform clonal frequency distribution. We determined evenness profiles for each repertoire and evaluated cross-repertoire evenness similarity by Pearson correlation of the repertoires’ evenness profiles as described previously (Amoriello et al., 2020; Greiff et al., 2015b, 2017a).

#### immuneREF Feature: Positional amino acid frequencies

The positional amino acid frequencies were calculated separately for each CDR3 sequence length. To decrease bias by extraordinarily short or long CDR3 sequences, we limited this analysis to a range of the most common lengths (8–20 amino acids) (Greiff et al., 2017a; Raybould et al., 2019). Briefly, per position amino acid frequencies are calculated for each length. Subsequently, the resulting per length frequency vectors of each repertoire were Pearson-correlated by length and the mean correlation was calculated. Unlike in the case of k-mer occurrences, no positions are excluded, making AA frequency more sensitive to VDJ usage perturbations. Relative frequencies were used for all positional amino acid frequency calculations.

#### immuneREF Feature: Sequence similarity network architecture

As previously described (Miho et al., 2019), we constructed a sequence similarity network for each immune repertoire: nodes represent amino acid CDR3 sequences connected by similarity edges if they had a Levenshtein Distance of 1 (LD=1). The igraph R package was used to calculate networks (v.1.2.4.1, Csardi and Nepusz, 2006), that were analyzed with respect to four measures representing different aspects of network architecture: (i) cumulative degree distribution, (ii) mean hub score (Kleinberg hub centrality score), (iii) fraction of unconnected clusters and nodes and (iv) percent of sequences in the largest connected component. An LD = 1 network was constructed for each repertoire and the similarity between the repertoires’ resulting network was evaluated with respect to their differences in the cumulative degree distribution, mean hub-score, outlier sequence occurrence, and largest network components; these metrics have been shown to be defining repertoire characteristics that are robust to subsampling (Miho et al., 2019).

The similarity of the architecture between two repertoires A and B was calculated as the mean of four components: (i) the cumulative degree distribution (Pearson correlation between repertoires), (ii) mean hub scores (1 – |*MeanHubScore_A_* – *MeanHubScore_B_*|), (iii) the fraction of unconnected components, and (iv) the fraction of sequences in the largest component (1 – |*PercLargestComponents_A_* – *PercLargestComponents_B_*|). Unlike many of the other features, the network feature combines multiple single measures, which rendered it difficult to perform Pearson correlation analysis involving all four investigated network measures. Therefore, we adopted the network feature comparison approach described above.

#### immuneREF Feature: Repertoire overlap (Convergence)

The pairwise repertoire clonal overlap (clones defined based on 100% similarity of CDR3 amino acid sequence), was calculated across repertoires, as previously described (Greiff et al., 2017b):

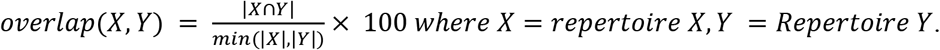

This clonal sequence overlap measure represents the similarity value between repertoires with respect to clonal convergence.

#### immuneREF Feature: Germline gene diversity

The relative frequency of germline genes (defined by the ImmunoGenetics Database, IMGT) (Giudicelli et al., 2004) across clones in each repertoire was calculated for each repertoire depending on species and immune receptor class (Ig, TR). The germline gene usage allows insight into deviations from a baseline recombinational likelihood and thereby captures the potential impact of disease, vaccine, or other events on the immune state (Avnir et al., 2016; Greiff et al., 2017a). To determine germline gene usage similarities, we examined the V- and J-gene frequencies across clones for each individual. The Pearson correlation coefficient was determined for each of the frequency vectors (V-, D-, J-gene) with entries of all IMGT variants in a pairwise fashion between samples as described previously (Greiff et al., 2017a; Weber et al., 2019). Specifically, the correlations are calculated per germline gene, leading to separate V_cor, D_cor, J_cor values (and additionally VJ_cor for each V_J combination). The resulting correlation values are combined into a single value by calculating a weighted mean of these components. The weight vector used for the results in the manuscript is c(V=1,D=1,J=1,VJ=0).

#### immuneREF Feature: Gapped k-mer occurrence

For a given k-mer size *k* and maximal gap length *m*, the nucleotide-based gapped-pair-k-mer occurrences were counted for all *gap sizes* ≤ *m* (Palme et al., 2015). The parameters *k* and *m* were chosen based on previous research (Greiff et al., 2017b), where defining parameters k = 3, m ≤ 3 was shown to lead to an encoding sufficient for sequence classification. The counts were normalized by the total number of gapped k-mers found across all gap sizes such that short-gap gapped-k-mers were weighted higher than larger gap sizes. While the amino acid frequency distribution contains positional information, the gapped k-mer occurrence represents short- and long-range sequential information encoded in the repertoire. We counted the occurrence of gapped k-mers (k = 3, m ≤ 3) across all CDR3 sequences of a repertoire and correlated the resulting distributions between repertoire pairs using Pearson correlation as described previously (Weber et al., 2019).

#### immuneREF Feature: transcriptome integration

In order to keep the most informative genes from the genes obtained in a transcriptome experiment, immuneREF firstly applies a low variation filter (Hackstadt and Hess, 2009). Specifically, the standard deviation (SD) is calculated per gene across samples, and all genes above a certain threshold (default, SD>1) are preserved for subsequent analysis.

To construct the gene expression feature similarity matrix, the Pearson correlation was calculated between samples. Additional approaches for the calculation of the gene expression feature similarity matrix implemented in the immuneREF package (mutual rank, PCA) are described in the package documentation.

### Calculating repertoire similarities per feature

The calculation of the similarity values between a pair of repertoires was performed in a feature-specific manner as described in the methods section of each feature.

### Repertoire similarity – Condensing features into a composite network

The single-features are condensed into a multi-feature network by taking the mean of all single-feature similarity values resulting in a single repertoire similarity value. The resulting condensed network represents a weighted composite of the single-feature similarity networks. Additional approaches to obtain a composite network (max similarity, min similarity, SNF (Wang et al., 2014)) are implemented in the R-package as described in the package documentation

### Mutual information

Mutual information is a measure that quantifies to what extent one random variable explains another. Mutual information was defined as

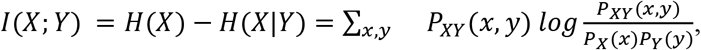

Where, H(X) is the marginal entropy, H(X|Y) the conditional entropy P_XY_ the joint probability distribution of X and Y and P_X_ and P_Y_ the respective marginals. Mutual information was calculated using the R packages entropy (v.1.2.1, Hausser, 2014) and infotheo (v.1.2.0, Meyer, 2014). The values were normalized to the range [0,1] by dividing the mutual information by the sum of the entropies H(X)+H(Y). This normalized mutual information, also known as redundancy, is zero when both are independent and maximal when knowledge of one of the variables becomes redundant given the other.

### Quantification of mutual information across ensembles of repertoire features

The mutual information between two features was calculated across all values in the similarity matrix, whereas the similarity matrix represents all pairwise similarity values between repertoires for a given feature. For the V(D)J diversity feature, values were set to zero by definition (i.e., the similarity between repertoires of different species/receptors) and were excluded from this calculation.

We ensembled immune information captured by the repertoire features (Fig. 2) as the extent to which repertoire features collectively cover immune repertoire complexity. Specifically, we evaluated the change in mutual information between subsequently added features. Features were added one by one (*1*-feature network → *2*-feature network, *2*-feature network → *3*-feature network and so forth, where *n*-feature means *n* features combined into a composite network), with the next feature to be chosen randomly (500 permutations of feature combinations per “n-features → *n*+*1*-feature” step).

**Figure 2.**
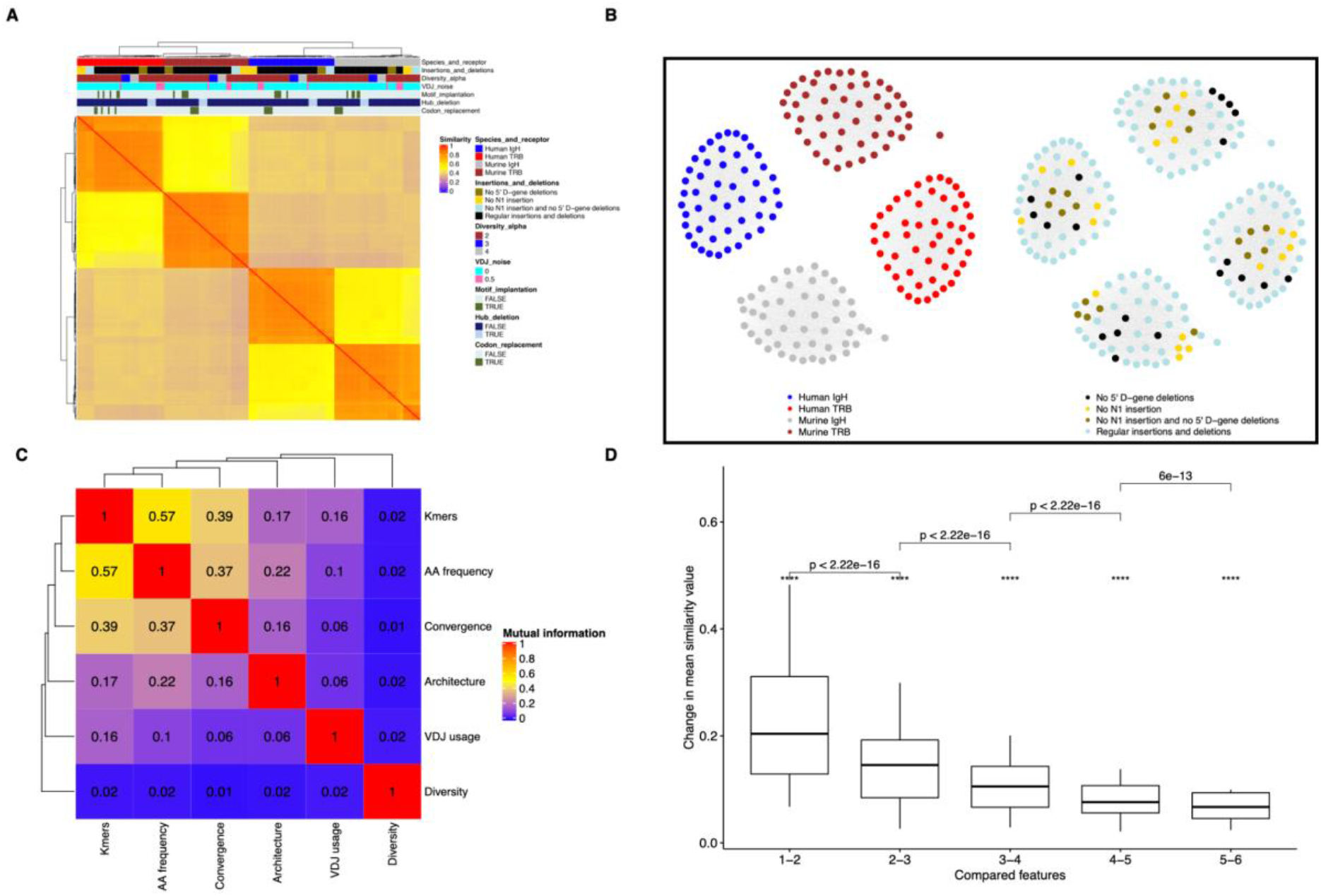
immuneREF measures immune repertoire similarity with high sensitivity using features that capture immune repertoire biology. We simulated 200 immune repertoires using 40 different parameter combinations (in quintuplicate). **(A)** Hierarchical clustering visualizes the sensitivity of immuneREF by the successful grouping of immune repertoires that were simulated with slightly different parameters (composite network, see main text for details). **(B)** Network visualization with simulated repertoires as nodes and weighted edges between repertoires of similarity values above the upper quartile. **(C)** Quantification of mutual information among immune repertoire features. **(D)** Change in mean similarity of composite networks of increasing number of features.

### Local repertoire similarity

To determine a single value measure for how connected a repertoire is within a subgraph (e.g. the repertoires of healthy human IgH repertoires and the similarity values between them), we defined the local similarity measure. It is calculated by dividing the node strength of each repertoire within a subgraph (sum of all edge weights connecting it to the other nodes in the subgraph) by the sum of all node strengths in the subgraph.

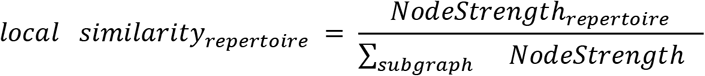

Local similarity gives the ratio of node strength that is connected to each repertoire in a subgraph and thus allows the identification of the most and least representative node of any category (the one most and least strongly connected within that category, respectively, see Figure 2C). Local similarity is dependent on the number of nodes within the subgraph and is therefore only used to compare repertoires within the subgraph. To enable comparison of local similarity values across different subgraphs, local similarity can be scaled by dividing by the number of nodes in the subgraph to correct for varying subgraph sizes in cases where the number of repertoires per subgraph differs.

### Simulation of adaptive immune receptor repertoires representing ground truth data

We simulated 200 immune repertoire datasets where we controlled 40 parameter combinations over multiple replicates, thus allowing us to generate datasets where there is *ground truth*. Simulated repertoires were generated by the immuneSIM framework (R package) (v.0.8.7, Weber et al., 2019). Each simulated repertoire contained 12’000 sequences and varied with respect to species (mouse, human), receptor (BCR, TCR), germline gene distribution, clone count distribution, the occurrence of N1, N2 insertions and deletions in V, D, and J genes. Additionally, a subset of repertoires was modified post-simulation: in order to simulate motif occurrence, the motifs “YAY” (“tacgcctac”) and “YVY” (“tacgtctac”) were implanted with a probability of 2.5% each at a random position in the complementarity determining region (CDR3). To create repertoires with variation in sequence similarity network architecture, the top 5% sequences with the highest hub scores in a given repertoire were removed. In order to evaluate the sensitivity of the gapped k-mer occurrence feature, repertoires that differ in nucleotide composition, while retaining amino acid composition, were generated by introducing synonymous codons (“tat” → “tac” for Tyrosine, “agt” → “agc” for Serine and “gtt” → “gtg” for Valine) in 50% of VDJ sequences. Finally, the simulated and modified repertoires were subsampled to 10’000 sequences to ensure equal dataset size. The simulation parameters and their expected impact on each feature are summarized in Supplementary Tables 1 and 2.

### Immune repertoire sequencing datasets

We conducted our analysis on 2’408 deep sequencing immune repertoires collected from four different studies: (i) a mouse immunization study of BCRs (flow cytometry-sorted B cells from different tissues: naive B cells from spleen (IgM), pre B cells (IgM) and IgG plasma cells from bone marrow, RNA-based high-throughput sequencing, preprocessed with MiXCR (Bolotin et al., 2015), for more details, please see (Greiff et al., 2017a)), (ii) a study of human TCRβ repertoires and signatures of cytomegalovirus, DNA-based high-throughput sequencing (CMV+/-, unsorted PBMC) (Emerson et al., 2017), (iii) a study of TCR repertoires of patients recovered from mild cases of Covid-19 (Minervina et al., 2021) and (iv) the PanImmune repertoire database (PIRD, unsorted PBMC, preprocessed with iMonitor) (Zhang et al., 2019) (see Supplementary Table 3). All sequences with stop codons were excluded and the naming of columns and V,D,J calls was standardized according to AIRR-community standards (Rubelt et al., 2017). When larger, each dataset was subsampled to 10’000 sequences (top clones by descending clonal frequency). Quality and read statistics may be found in the respective publications.

### Statistical analysis and graphics software

Statistical analysis was performed using R 3.6.1 (R Core Team). Graphics were generated using the R packages ggplot2 v3.2.1 (Wickham, 2009), ggbeeswarm v0.6.0 (Erik Clarke and Scott Sherrill-Mix, 2017), RColorBrewer v1.1-2 (Neuwirth, 2014), ComplexHeatmap v2.2.0 (heatmaps) (Gu, 2015), igraph v1.2.4.2 (network plots) (Csardi and Nepusz, 2006), ggiraphExtra v.0.2.9 (radar plots) (Keon-Woong Moon, 2018), GGally v1.4.0 (parallel plots) (Schloerke et al., 2018). Parallel computing immuneREF analysis was performed using the R packages foreach v1.4.7 (Microsoft and Steve Weston, 2019) and doMC v1.3.6 (Revolution Analytics and Steve Weston, 2019). Figure 1 was created using Biorender.com.

### immuneREF R-package and documentation

The immuneREF analysis workflow is made available via the immuneREF R package hosted on GitHub (https://github.com/GreiffLab/immuneREF). Documentation of the immuneREF package is provided on readthedocs (https://immuneref.readthedocs.io).

## Results

### Reference-based comparison of immune repertoires based on immunological features: constructing a similarity atlas of immune repertoires

To derive a similarity measure for immune repertoires, we devised a framework that calculates a repertoire similarity score based on six features that reflect immune repertoire biology (Fig. 1, Supplementary Table 4). These features are (1) germline gene diversity (Greiff et al., 2015c; Yaari and Kleinstein, 2015), (2) clonal diversity (Greiff et al., 2015b; Stern et al., 2014), (3) clonal overlap (Greiff et al., 2015c; Yaari and Kleinstein, 2015), (4) positional amino acid frequencies (Mason et al., 2019), (5) repertoire similarity architecture (Bashford-Rogers et al., 2013; Ben-Hamo and Efroni, 2011; Miho et al., 2019) and (6) k-mer occurrence (Greiff et al., 2017b; Thomas et al., 2014) (see the *Methods* Section for a detailed immunological and mathematical description of these features). A similarity score is calculated for each pair of repertoires and each feature (six *n* x *n* symmetric matrices, *n* = number of repertoires), creating a similarity matrix for each feature. This matrix may be viewed as a weighted network, in which the nodes correspond to repertoires and the edges connecting the nodes are the similarity scores. The resulting six single feature similarity networks enable insight into per-feature similarity. Finally, a composite network of the six feature similarity networks represents an interpretable multidimensional picture of the repertoire landscape. Briefly, the single-features are condensed into a multi-feature composite network by taking the mean of all single-feature similarity values resulting in a single repertoire similarity value (for alternative approaches to computing composite networks, see Methods section). By virtue of representing a similarity matrix as a weighted network repertoire, similarity may be computed on selected levels such as one(repertoire)-to-many (repertoires), many-to-one, and many-to-many (Fig. 1). Interpretability stems from all repertoire features being transformed into a similarity measure on a 0–1 scale allowing for direct quantification of their individual contribution to multidimensional immune repertoire similarity.

### immuneREF measures immune repertoire similarity with high sensitivity

We sought to quantify the sensitivity by which immuneREF can detect differences between immune repertoires with respect to the six repertoire features. The simulated repertoires, varying in a controlled manner, represent a ground truth reference map that enables a more precise assessment of immuneREF sensitivity. For example, simulated repertoires may be used to guide the evaluation of variation between experimental repertoires with respect to each repertoire feature as well as multi-feature combinations. Simulations were performed using the immuneSIM repertoire simulation suite (Weber et al., 2019), which was used to create native-like repertoires that were varied across eight parameters. Native-likeness was demonstrated in (Weber et al., 2019). The parameters that were varied across simulated repertoires included clone count distribution, V-, (D-), J-gene frequency noise, insertion and deletion likelihoods, species (human and mouse), and receptor type (IgH, TRB). We constructed additional simulated repertoires with spiked-in motifs (mimicking antigen-binding motifs (Akbar et al., 2019)), excluded hub sequences in the sequence similarity network (simulating network architecture variation (Miho et al., 2019)) and replaced nucleotide codons with synonymous codons (simulating biases in the k-mer occurrence that are relevant in detectable immunogenomic patterns of public clones (Greiff et al., 2017b)) (see *Methods*, Supplementary Table 1 lists the parameter variations used for the simulations, Supplementary Table 2 summarizes how each of the parameters is expected to influence the six immuneREF features). The parameter combinations were chosen such that each simulated repertoire varied only along one parameter dimension at a time, allowing us to determine the sensitivity of each feature to each parameter change.

The mathematical structure of the single-feature similarity matrices enables their merging into a composite network that provides the opportunity for a condensed single-score representation of inter-sample repertoire similarity. The composite immuneREF network (which combines all six repertoire features) recovers major variation in the repertoires including noise introducing parameter changes (Figs. 2AB). immuneREF also clearly distinguishes repertoires from different receptors and species based on strongly distinguishing features such as V-, (D-), J-gene usage while allowing the identification of commonalities in amino acid usage, clonal diversity, and architecture across immune receptors and species. This sensitivity analysis also underlines a major advantage of immuneREF, namely its flexibility to accommodate both BCR and TCR repertoires from different species in one single analysis workflow.

We quantified the sensitivity of immuneREF by detecting significant changes in similarity scores corresponding to the variation in simulation parameters across both the single feature (Supplementary Figs. 2–4) and composite network (Fig. 2A) and found that each feature had a unique sensitivity profile to changes in the simulation parameters, underscoring the value of per-feature similarity evaluation. For example, a change in the alpha parameter of the Hill function (controlling clone count distribution) solely impacted the immuneREF diversity feature. As the immuneSIM parameter controlling the distribution of clone counts only affects the clone count simulation without impacting simulated sequences, the fact that only the feature targeted by the parameter change is impacted shows that immuneREF is robust to random noise in the simulation that is not introduced through parameter changes. An increase in the V(D)J noise parameter, which modifies the frequencies of the germline genes used in the simulation, led to detectable and significant changes in similarities of the germline gene usage and k-mer occurrence features. Modification of the insertion/deletion patterns (dropout of deletions and or insertions) led to a consistent impact in the amino acid frequency feature and, more importantly, the architecture feature, where a lower diversity due to restricted insertions and deletions led to significant changes in network architecture. Implanting motifs at various frequencies led to a significant similarity change in the k-mer occurrence feature. The deletion of hub sequences led to an impact in the architecture feature and also changed the repertoire overlap similarity, thus underlining the importance of public clones in the network architecture as reported previously (Miho et al., 2019). Finally, we modified the repertoires by introducing synonymous codons at various percentages and found that the k-mer occurrence feature was the only one impacted. Therefore, we conclude that immuneREF features largely react as hypothesized to variation in simulation parameters (Supplementary Table 2). Taken together, we demonstrated that the immuneREF framework is sensitive to even comparatively small repertoire variations.

### Mutual information analysis demonstrates no to limited inter-dependence of immuneREF features

While the examined features were initially chosen based on immunological criteria, we also wished to verify whether each feature provides a sufficiently different measurement of the immune repertoire information space Fig. 2C). Specifically, having integrated all features into a common coordinate system, we were able to compute cross-feature mutual information and found that features show no to limited dependence (range=0.01–0.57, Fig. 2C) indicating largely non-overlapping and distinct spaces of immune information captured. The highest mutual information was found between the positional and sequential sequence-derived features (i.e., positional amino acid frequency and gapped k-mer occurrence, respectively) while the lowest mutual information value was found between the diversity and convergence features (Fig. 2C).

Complementarily, we sought to quantify to what extent the addition of new repertoire features leads to diminishing returns (sufficiency analysis). To this end, we computed the mean change in repertoire similarity values when increasing the number of features from 1 through 6. Thereby, we could show that each additional feature added increasingly less information, as shown by the diminishing change of the mean similarity value with each added feature. The saturation of the mean similarity change curve indicated information saturation independent of the order in which features were arranged (Fig. 2D, Supplementary Fig. 5). As discussed below, mutual information values behaved similarly for experimental repertoire data. Thus, we demonstrated that the immuneREF framework creates information-laden similarity networks, whose topologies capture the immunological similarity landscape of immune repertoires.

### The similarity landscape of simulated repertoires defines reference repertoires

By calculating the similarity matrix for each of the six immune repertoire features, we embedded the six different immunological features into a common coordinate system, i.e., a network structure. This network (with nodes representing repertoires and weighted edges representing pairwise similarity) situates each repertoire within a similarity landscape allowing to quantify many-to-many repertoire similarity.

A more fine-grained image of the similarity landscape may be gained by examining the similarity from the perspective of every single repertoire (Figs. 3C,D). We define the local similarity of a repertoire to its neighboring repertoires as a scaled node strength (see Methods). This local similarity represents the position of the repertoire with respect to its direct neighbors in its cohort (defined by an application-dependent label, e.g., same species and disease) and allows us to distinguish between well embedded and aberrant repertoires. The local similarity measure further acts as a magnifying glass by elucidating finer differences between repertoires, which are diluted by population averages when examining repertoire similarity across the full similarity network. Using this perspective, repertoires that are most (locally) similar to other repertoires in their cohort can be identified, allowing the extraction of repertoires most representative for a given immune state. Such detailed one-to-one feature comparisons highlight, in the most simple case, which features of the simulated repertoires are receptor-specific (amino acid frequency, k-mer occurrence, VDJ usage, and convergence) and which are more general to immune repertoire data showing higher similarity across different species and receptors (diversity, architecture) (Fig. 3E, Supplementary Fig. 6).

**Figure 3.**
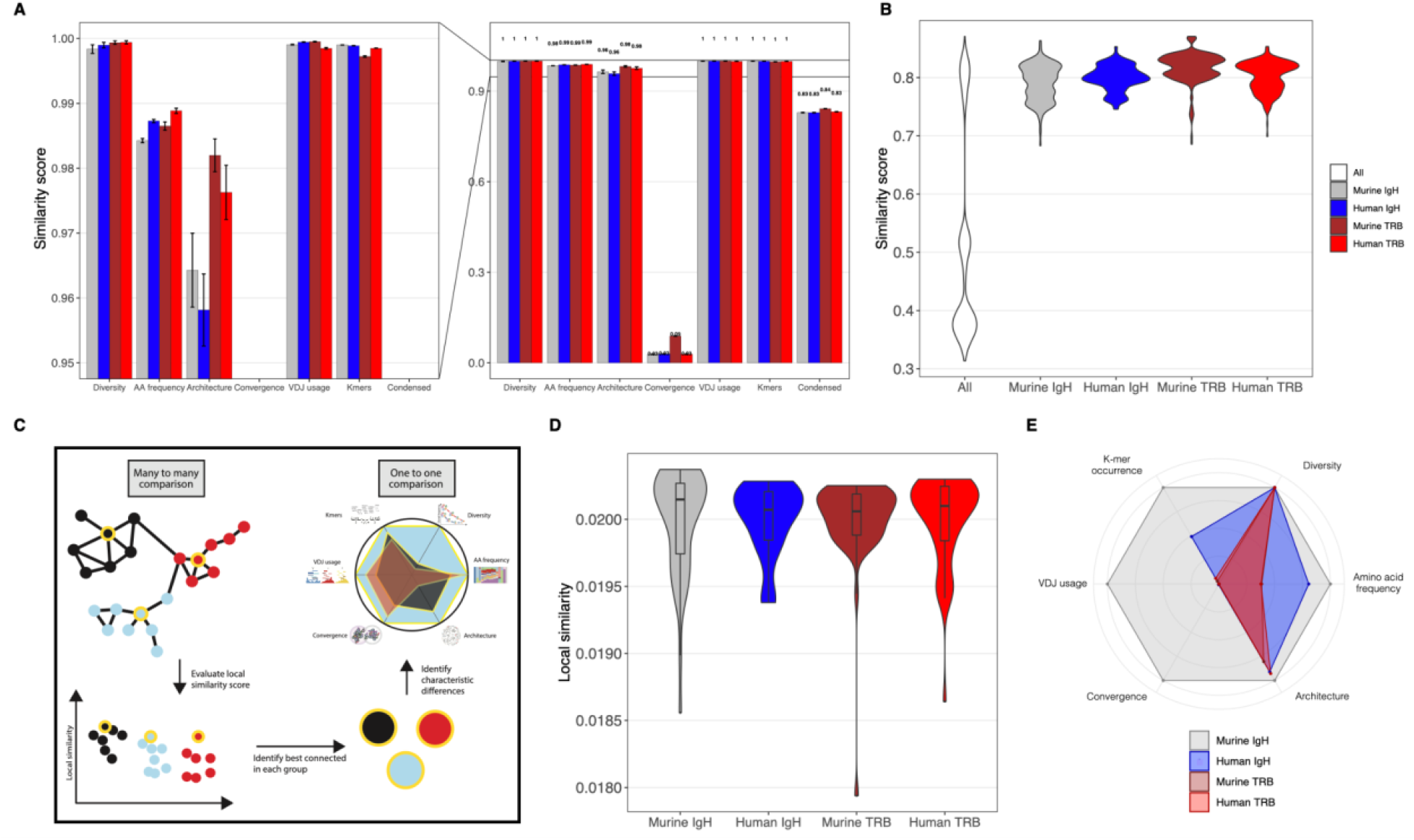
The similarity landscape of simulated repertoires defines reference repertoires. **(A)** Baseline similarity between replicates for repertoires simulated using default immuneSIM parameters (see Supplementary Table 1) is ≥0.96 for five of six features, the convergence feature being the exception by definition at ≤0.09. **(B)** Repertoire similarity distribution in a condensed network across the various evaluated parameter range. Across cohorts, similarity scores have a broad range, while within cohorts the range is more restricted. **(C)** Workflow to determine representative repertoires per cohort going from many-to-many to a one-to-one comparison. **(D)** Local similarity distribution per species/receptor combination enables situating each repertoire based on its connectivity with respect to neighbors in the same cohort. **(E)** Comparing repertoires with maximal local similarity in their cohort visualizes the commonalities between receptor types, here the Murine IgH repertoire with maximal local similarity serves as a reference repertoire. The plot visualizes the similarities of each non-reference repertoire to the Murine IgH reference.

Having evaluated the similarity of simulated datasets, these may serve as a reference to interpreting similarity score variation of experimental repertoires (Fig. 3C), thus enabling the creation of equivalence classes of immune repertoires not only as previously performed based on clonal expansion (Greiff Genome Medicine 2015) but based on six repertoire features. Furthermore, any evaluated repertoire, be it of experimental or simulation origin, will become a new node in the similarity network and may serve as a valid reference point (just as any other node in the network). This network of self-augmenting repertoire similarity reference points is another source of interpretability as it allows the linking of the repertoire similarity of any number of repertoires with their underlying features. In the next section, we provide such a repertoire similarity network on experimental datasets.

### Validation of immuneREF on experimental data: detection of differences between cell populations in mouse immunization and human Covid-19 datasets

To validate immuneREF sensitivity on experimental data, we used antibody repertoire datasets generated from a mouse antigen immunization study, where differences in the similarity between antigen immunization cohorts are expected (Greiff et al., 2017b, 2017a; Miho et al., 2019). Notably, we were able to recover clear differences between isotypes and cell populations (both with higher within-cohort and lower across-cohort similarity), and additionally found that the antigen immunization cohorts have more distinct similarity profiles in the plasma cell populations (IgG) compared to the antigen-inexperienced cell populations (Supplementary Figs. 7, 8). The overall high similarity scores across the full immunological feature range are in agreement with our previous studies where we observed high similarity between these repertoires on a single feature basis (Greiff et al., 2017a).

Similarly, applying immuneREF to TCR repertoires of patients recovered from mild cases of Covid-19 (Minervina et al., 2021), revealed clusters of increased similarity within patients and cell populations (Supplementary Fig. 9).

### Application of immuneREF to >1500 experimental blood immune repertoires indicates only small similarity-based differences between health and autoimmune disease

Having established the sensitivity of our approach in detecting a wide range of differences between simulated repertoires (Figs. 2,3) and between experimental repertoires of different B cell populations (Supplementary Figs. 7–9) with respect to immunologically relevant and interpretable repertoire features, we set out to determine the similarity landscape of large scale experimental TCR repertoire datasets. We evaluated 1,522 human TCR repertoires derived from peripheral blood mononuclear cells (PBMCs) of patients with varying and diverse immune states (PanImmune Repertoire Database (PIRD) dataset containing samples from Healthy, Rheumatoid Arthritis (RA) and Systemic Lupus Erythematosus (SLE) patients, Supplementary Table 3). We found an even similarity landscape of overall high similarity scores (Fig. 4A). Similarity score distribution was also even in single features, which despite feature-specific differences, show overall high similarity scores between repertoires. We examined networks at three different similarity cutoffs (an edge is drawn between two repertoire nodes if their similarity is in 25, 50, and 75% top weights, respectively), and found that in all three cases no immune state-specific grouping could be observed (Fig. 4B).

**Figure 4.**
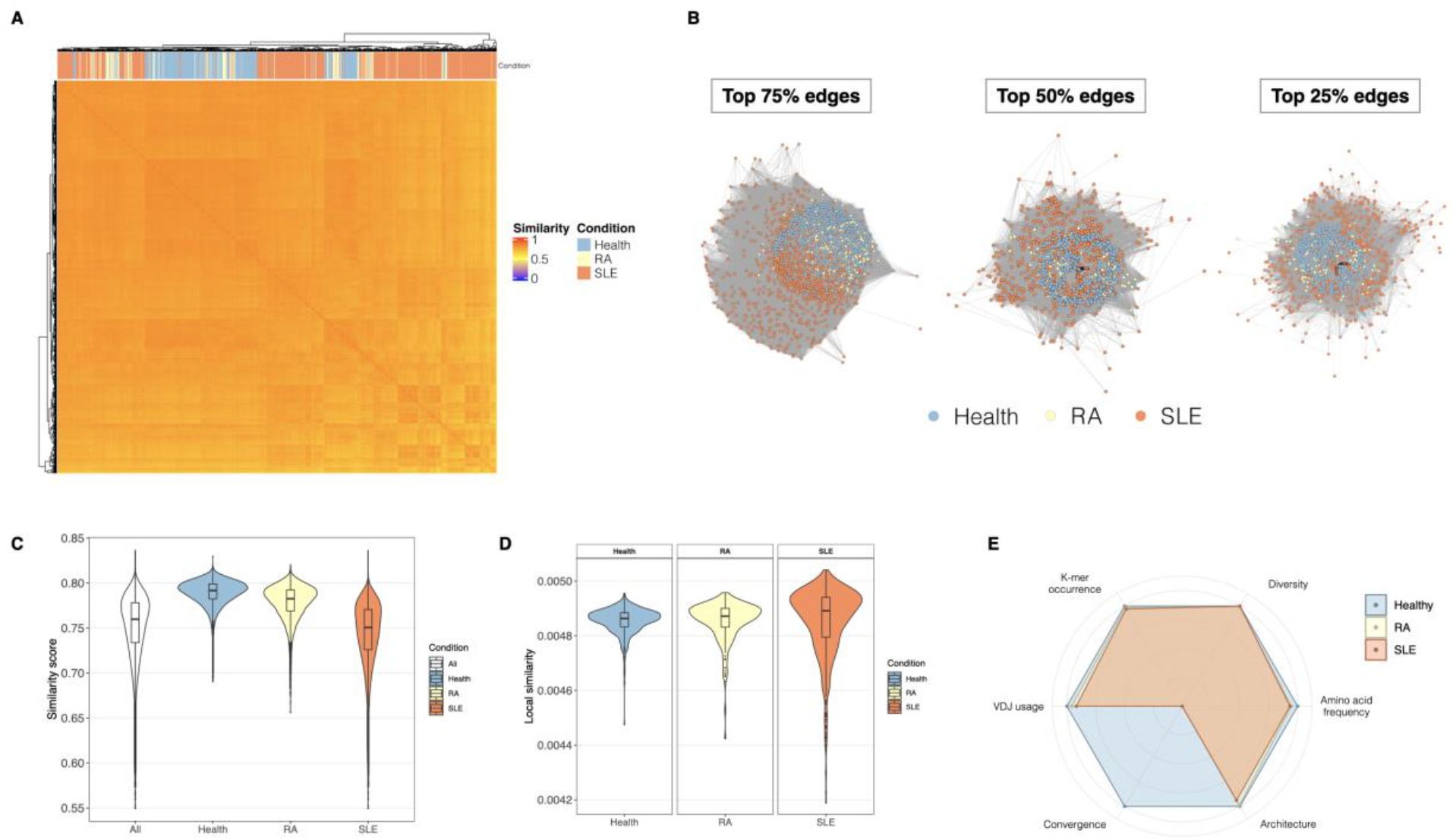
Application of immuneREF to 1,522 experimental repertoires. **(A)** Similarity landscape of experimental (human, TCR) repertoires across three immune states (Healthy: 439 repertoires, Rheumatoid Arthritis: 206 repertoires and Systemic Lupus Erythematosus: 877 repertoires) **(B)** Network visualization of the 1,721 nodes and weighted edges between repertoires of similarity scores (at three cutoff levels, 25%, 50% and 75% top edge weights). **(C)** Distribution of similarity score across the entire network and per immune state shows different degrees of within cohort homogeneity. **(D)** Distribution of local similarity values per repertoire, faceted by cohort. **(E)** Comparison of the repertoires with the highest local similarity per immune state and an immuneSIM reference repertoire (default immuneSIM parameters, see Supplementary Table 1).

The range of general and local similarities across all samples as well as within each disease cohort was evaluated using an analogous approach to that used for the simulated datasets (Figs. 4CD). While the similarity scores ranged between ~0.5–0.8 overall, the within-disease cohort spread varied, with the Healthy and RA cohorts showing a more restricted range of similarity scores compared to a broader range for SLE (Fig. 4CD).

To quantify per feature similarity and dissimilarity with respect to a reference dataset, we compared the repertoires identified as the ones best-connected (highest local similarity) within their cohort, to an immuneSIM reference repertoire (human, TRB, standard parameters, see Methods) (Fig. 4E). The similarity scores of all tested immune states largely overlap with respect to the healthy reference repertoire, with convergence being the feature dimension with the largest dissimilarity, meaning there is almost no convergence between the RA or SLE samples and the reference. Comparing the single features of the so determined most and least ‘representative’ repertoires per immune state (Supplementary Fig. 10) further underlines the high similarity between repertoires of identical and different immune states.

Following our observations of high repertoire similarity within the PIRD dataset, we ran immuneREF on another large publicly available dataset (human, TCR) (Emerson et al., 2017) with yet another difference in immune state (CMV). The dataset contains 666 PBMC samples of which 289 are from CMV positive patients, 351 are from CMV negative patients and 26 are from patients with unknown CMV status. This dataset has previously been used to showcase immune state classification with high accuracy via the identification of CMV-associated public TCR sequences (sequences shared between individuals). In a similar fashion, immune state-associated public sequences were used to successfully classify RA and SLE samples from the PIRD dataset (Liu et al., 2019). As with the PIRD dataset, we observed high within and across immune state repertoire similarity (Supplementary Figs. 11, 12). This is in line with the findings of Emerson and colleagues as they found that only a small subset of clones (CMV-associated in Emerson et al., 2017) significantly differed in abundance between immune states (CMV+, CMV-) and that that shared antigen exposure to CMV led to a reduced number of shared TCRβ clones, even after controlling for individual HLA type, indicating a largely private response to a major viral antigenic exposure (Johnson et al., 2021).

In summary, the results of our analysis of human TCR repertoires strongly support the argument that the signal-to-noise ratios, where signal means repertoire features associated with disease status, is unfavorably tilted towards noise, where noise is defined as technological and immunological information, which cannot *yet* be linked to a given disease state.

### Extensibility of immuneREF: Integration of gene expression with immune repertoire data

The mathematical structure of the composite network obtained from immuneREF allows the extensibility of the immuneREF framework to other features. As a proof of principle of this immuneREF capability, we show here an integrative analysis of immune repertoires and gene expression. This integration is of high interest to RNA-seq experiments that include both receptor and global transcript sequences, or even repertoire experiments paired with transcriptomics (Song et al., 2021). Integration of immune repertoire with gene expression is challenging due to the multidimensional nature of both kinds of datasets and the discrepancy in their data structure. Current attempts of integration are still over-simplistic, such as the calculation of correlation between the number of distinct CDR3 amino acid sequences and gene expression of some marker genes such as *CD3*, *CD4*, *CD8*, HLA Class I and Class II genes (Brown et al. Genome Medicine, 2015, 7:125).

immuneREF includes the option to evaluate similarity based on a gene expression matrix and add it to the composite network. Briefly, immuneREF first filters all genes with low variation between experimental conditions and then calculates the pairwise correlation between observations to construct a single gene expression feature (similarity matrix). Once the seven features (six from immune repertoires and one for gene expression) are calculated, they may be condensed into a multi-feature network as described above. Our solution for integrating receptors with gene expression confers immuneREF the advantage of overlaying dual biological information (Supplementary Fig. 13A).

As an example, we analyzed bulk RNA-seq gene expression of pre-B-cell line B3 from the published STATegra project (Gomez-Cabrero et al. Sci Data, 2019, 6: 256). This is a time-course experiment that collects samples at six time points using an inducible Ikaros system where B cell progenitors undergo growth arrest and differentiation (Supplementary Fig. 13B). PCA showed clear differences at gene expression level when control and Ikaros groups were compared but also within the Ikaros group across time, being t0 the nearest to controls (Supplementary Fig. 13B). To generate the single feature similarity matrix of gene expression that better collects these differences, we tested the 3 available correlation-based methods implemented in immuneREF (Supplementary Fig. 13C–E). All of them perfectly separated control (blue) and Ikaros (red) groups. Additionally, “Pearson correlation” and “PCA scores” nearly recovered correctly the time series pattern (purple to yellow degradation), while Mutual rank matched perfectly.

## Discussion

Combining methods from both immune repertoire and network analysis, we have provided a framework for flexible reference-based quantification of immune repertoire similarity. Using ground truth simulations, we show that immuneREF is sensitive to inter-repertoire differences in all immunological features. Taking advantage of information theory, we showed using both simulated and experimental data that the features selected for immuneREF cover a large extent of immune repertoire biology. We introduced the concepts of full-network repertoire similarity and local similarity, which allow complementary quantification of the impact of the differences in the repertoire similarity landscape. Specifically, while the more general repertoire similarity evaluated on the entire network provides insight into the range of similarity within and across conditions, local similarity shows a particular advantage of the network approach, as the embedding of a repertoire in its neighborhood can markedly differ from what can be expected by its pairwise connections.

immuneREF not only provides a framework for measuring immune repertoire similarity but also for interpreting it (Supplementary Fig.1). Specifically, it enables the creation of equivalence classes of immune repertoires lacking from existing methods (Supplementary Table 4). For example, once the similarity observed within a given set of experimentally obtained immune repertoires has been computed, such repertoires may function as reference points that in turn enable the interpretation of relative similarity in other repertoires (Figs. 1,3C). Of note, the concept of diversity measures creating equivalence classes has been noted previously for Hill diversity measures (Greiff et al., 2015b) and is here extended to include additional repertoire features immuneREF unifies single and composite features, frequency-dependent, and sequence-dependent similarity measures into one computational framework. Beyond quantifying the repertoire similarity of experimental immune repertoires, immuneREF also enables the comparison of simulated (Han et al., 2021; Marcou et al., 2017; Safonova et al., 2015; Weber et al., 2020)(Marcou et al., 2017; Safonova et al., 2015; Weber et al., 2019) and in vitro synthetic immune repertoires used for therapeutic antibody discovery (Mason et al., 2018). Furthermore, immuneREF may be used for data curation purposes in immune repertoire databases such as iReceptor (Corrie et al., 2018), VDJserver (Cowell et al., 2015), PIRD (Zhang et al., 2019), and Observed Antibody Space (Kovaltsuk et al., 2018). Specifically, upon the integration of an immune repertoire into a database, the similarity of the repertoire with all other stored repertoires may be computed. Beyond immunological insight, immuneREF may reveal unexpected variation, thus motivating follow-up inspection. Since immuneREF has been built to work across species, cell populations, and receptor types, experimental or simulated data (all-in-one comparative framework), it enables rapid distinction of cohort-specific and cohort-unspecific features. This is also important for comparative immunological approaches not centered on health vs. disease comparison but, for example, the evolution of adaptive immunity (Pancer and Cooper, 2006).

While we consider the usefulness of the six chosen features to be established (Fig. 2, Supplementary Fig. 5), we concede that the asymptotic nature of the sufficiency calculation leaves the door open to the introduction of additional features. The proposed set of immuneREF features denotes in this sense a minimally sufficient set for the analysis of immune repertoire datasets. It ensures sufficient coverage of the major variation-introducing aspects. It is for that reason we devised immuneREF as inherently modular, allowing single and multi-feature analysis as well as encouraging the addition of new features relevant for particular problems such as transcriptome analysis (Supplementary Fig. 13), HLA typing for TCR studies (DeWitt et al., 2018; Emerson et al., 2017; Francis et al., 2021), single-cell omics information (Han et al., 2021; Setliff et al., 2019; Sturm et al., 2020; Yermanos et al., 2021), gene-specific substitution profiles for somatic hypermutation analysis (Sheng et al., 2017), lineage-specific information (Hoehn et al., 2016, 2021), and antigen-specific and antigen-associated motifs identified by sequence clustering and machine learning (Akbar et al., 2019; Dash et al., 2017; Friedensohn et al., 2020; Glanville et al., 2017; Greiff et al., 2017b; Horst et al., 2021; Mason et al., 2019; Mayer-Blackwell et al., 2021; Meysman et al., 2018; Sidhom et al., 2019; Wong et al., 2020; Yohannes et al., 2021). In particular, a future extension of immuneREF may be a feature that reliably identifies antigen-specific sequences thus increasing the amount of immune information recovered. More generally, adult repertoires are very complex and contain hidden information of many antigens at different time points that might have been shared by different individuals. For instance, repertoire fingerprints of influenza infection might be present on most studied individuals and could explain the difficulty to distinguish healthy and diseased individuals. New features including (single-cell-based) antigen specificity patterns may help separate shared infection marks on the immune repertoire.

The ease of use of the immuneREF approach opens up new possibilities for large-scale comparative studies as shown on the PIRD dataset, which may yield additional insight into the challenges of predicting immune state based on repertoire profiling. Indeed, we found that the population average quantified by immuneREF may ‘‘conceal’’ relevant immunological phenotype signals – despite the fact that the sensitivity of immuneREF was shown to be high in simulated and experimental data (Fig. 2, Supplementary Figs. 7, 9). Given the lack of large-scale (antigen-specific) data, it remains unclear how the information of the immune state is distributed across immunological features. Specifically, our finding – that repertoire similarity does not differ across immune states – is strictly only valid for unsorted PBMC TCR repertoire data as examined in this study. As known from previous studies (Amoriello et al., 2020, 2021; Csepregi et al., 2021; Ghraichy et al., 2021; Greiff et al., 2017b; Li et al., 2020; Ota et al., 2022; Riedel et al., 2020; Rosati et al., 2021)r, different cell populations (in different lymphoid organs) may behave in a highly different manner (Supplementary Figs. 7, 9). On the other hand, it did not escape our attention that this broad similarity in human blood samples might suggest the maintenance of lymphocyte homeostasis even in the event of chronic disease.

Our results reinforce the notion that while some diseases may introduce abnormalities into the immune repertoire, others result in a comparatively normal one (Bashford-Rogers et al., 2019), a result that suggests the absence of a signature unique to health. If this is true, then blood-based immune repertoire diagnostics will require even more advanced methodologies to be developed (Widrich et al., 2020c). For example, for simulated repertoires, motif implants in ≥ 10% of sequences were required to affect the amino acid frequency and architecture features, suggesting that even in the case of high clonal expansion, the impact on the repertoire might not be sufficient to significantly change major repertoire features. This is reinforced by results showing that the disease-driving response in multiple autoimmune diseases is only to a small part antigen-specific (Christophersen et al., 2019). More generally, our paper advances the state-of-the-art of the immune repertoire field by changing the null hypothesis. Specifically, currently, the predominant thinking is that any immune state changes measurably the immune repertoire in a systematic fashion. Our paper challenges this view by finding that, *a priori*, we should not expect to see differences (Fig. 4, S10) and any substantial change must be proven. This change of perspective is highly valuable to the field as it pushes it towards more sensitive and robust approaches to immune repertoire and machine learning analysis (Kanduri et al., 2021; Slabodkin et al., 2021). Specifically, the usefulness of global features for diagnostics is severely limited and to detect single sequence-level differences (Emerson et al., 2017; Kanduri et al., 2021; Widrich et al., 2020a), single-sequence level statistical and machine learning approaches are needed (Greiff et al., 2020; Schattgen et al., 2021).

In the future, ultra-deep (Briney et al., 2019; Soto et al., 2019, 2020) and population-wide large-scale immune repertoire projects such as Human Vaccines Project (Crowe Jr. and Koff, 2015) may benefit from using immuneREF for identifying immune event-driven aberrations from a baseline repertoire similarity. Furthermore, large-scale database initiatives such as the iReceptor gateway (Corrie et al., 2018) may benefit from immuneREF functionality for on-the-fly computation of inter-dataset similarity.

## Supplementary Tables and Figures

**Supplementary Table 1.**
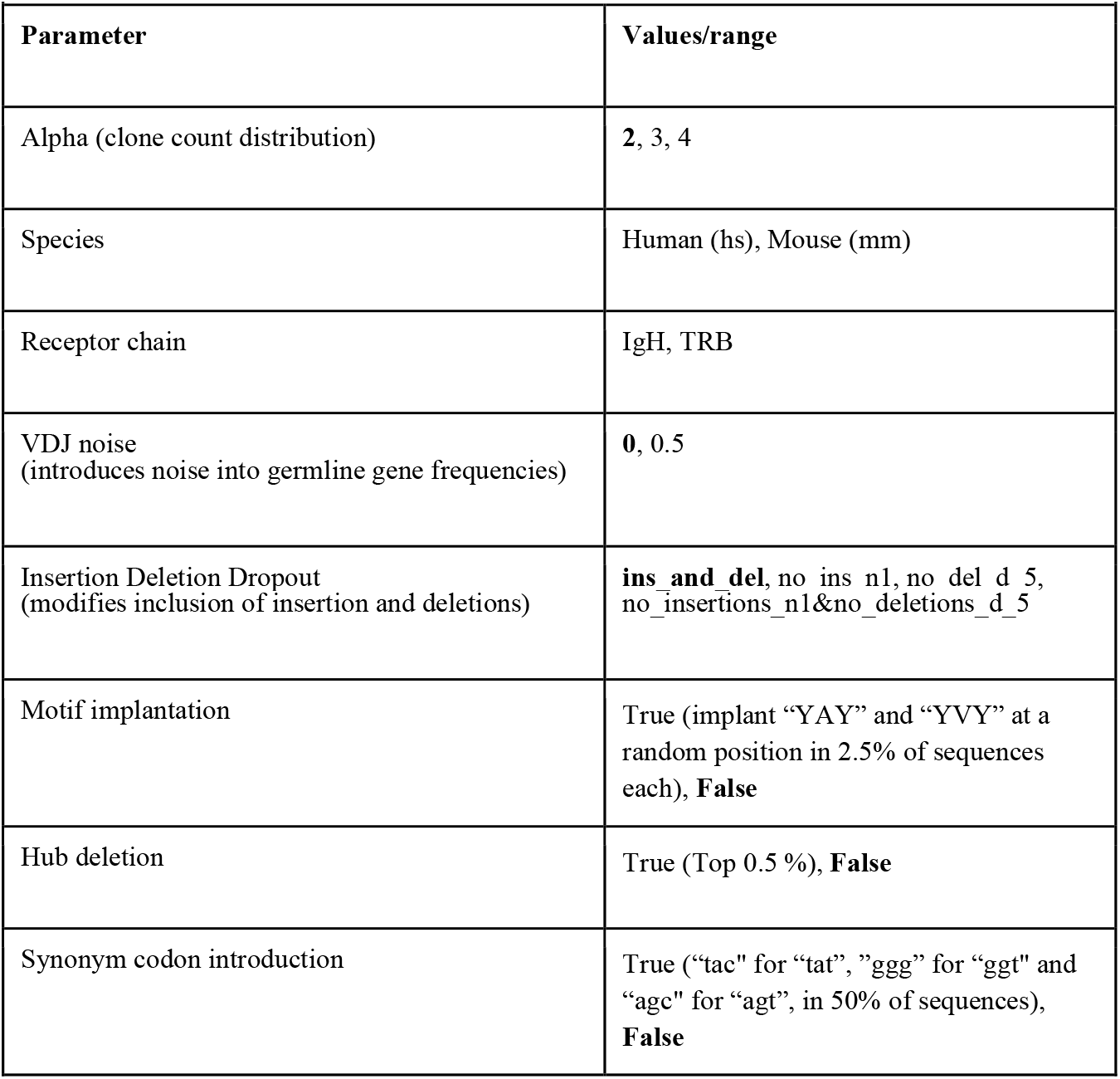
ImmuneSIM parameters for simulated datasets. The repertoires were simulated to differ across eight parameters. (In bold: Default parameters.) Please refer to immuneSIM documentation (https://immunesim.readthedocs.io/en/latest/parameters.html) for parameter definitions.

**Supplementary Table 2.** Hypothesized impact of parameter variation on immuneREF features. Theorized impact of each parameter variation on immuneREF features.

| Supplementary Table 2. Hypothesized impact of parameter variation on immuneREF features. |  |
| --- | --- |
| Parameter | Hypothesized Impact (Feature) |
| Alpha (clone count distribution) | Diversity feature only |
| Species | All except diversity |
| Receptor chain | All except diversity |
| VDJ noise<br>(introduces noise into VDJ frequencies) | Germline gene usage, kmer, A.A. frequency,<br>Architecture |
| Insertion Deletion Dropout<br>(modifies inclusion of insertion and deletions) | Germline gene usage, kmer, A.A. frequency,<br>Architecture |
| Motif implantation | Convergence, (minor: architecture) |
| Hub deletion | Architecture, Convergence |
| Synonym codon introduction | K-mer, (minor: Convergence) |

**Supplementary Table 3.** Quantitative description of high-throughput sequencing datasets. Overview of the number of unique CDR3 sequences of the experimental datasets analyzed using immuneREF. Where the number of clonal sequences exceeded 10’000, the immune repertoire was subsampled by extracting the 10’000 clonal sequences with the highest clone count.

| **Supplementary Table 3. Quantitative description of high-throughput sequencing datasets.** |  |  |
| --- | --- | --- |
| **Data Origin** | **Cell Type** | **No. of unique CDR3 sequences**<br>**(mean±std.error)** |
| Mouse (C57BL/6J), B cell (Greiff et al., 2017a) | preBC (IGM) (19)<br>nBC (IGM) (19)<br>PC (IGM) (19)<br>PC (IGG) (19) | 172'262 ± 9'394<br>395'558 ± 18'341<br>28'762 ± 11'443<br>147 ± 18 |
| Human (CMV-dataset), T cell (Emerson et al., 2017) | PBMC (CMV positive) (289)<br>PBMC (CMV negative) (351)<br>PBMC (CMV unknown) (26) | 177'095 ± 4'235<br>186'951 ± 4'029<br>154'478 ± 14386 |
| Human (Covid-dataset), T cell (TCRbeta) (Minervina et al., 2021) | CD4 (mild Covid) (10)<br>CD8 (mild Covid) (10)<br>Mem (mild Covid) (28)<br>PBMC (mild Covid) (24) | 321'827 ± 34'100<br>189'289 ± 29'240<br>36'233 ± 6'232<br>623'003 ± 46'519 |
| Human (Covid-dataset), T cell (TCRalpha) (Minervina et al., 2021) | CD4 (mild Covid) (10)<br>CD8 (mild Covid) (10)<br>Mem (mild Covid) (28)<br>PBMC (mild Covid) (24) | 233'948 ± 32'085<br>141'119 ± 20'064<br>29'073 ± 5'253<br>425'560 ± 28'554 |
| Human (BGI), T cell (Zhang et al., 2019) | PBMC (Healthy) (439))<br>PBMC (RA) (206)<br>PBMC (SLE) (877) | 74'195 ± 25'364<br>123'292 ± 68'630<br>84'582 ± 52'944 |

**Supplementary Table 4.**
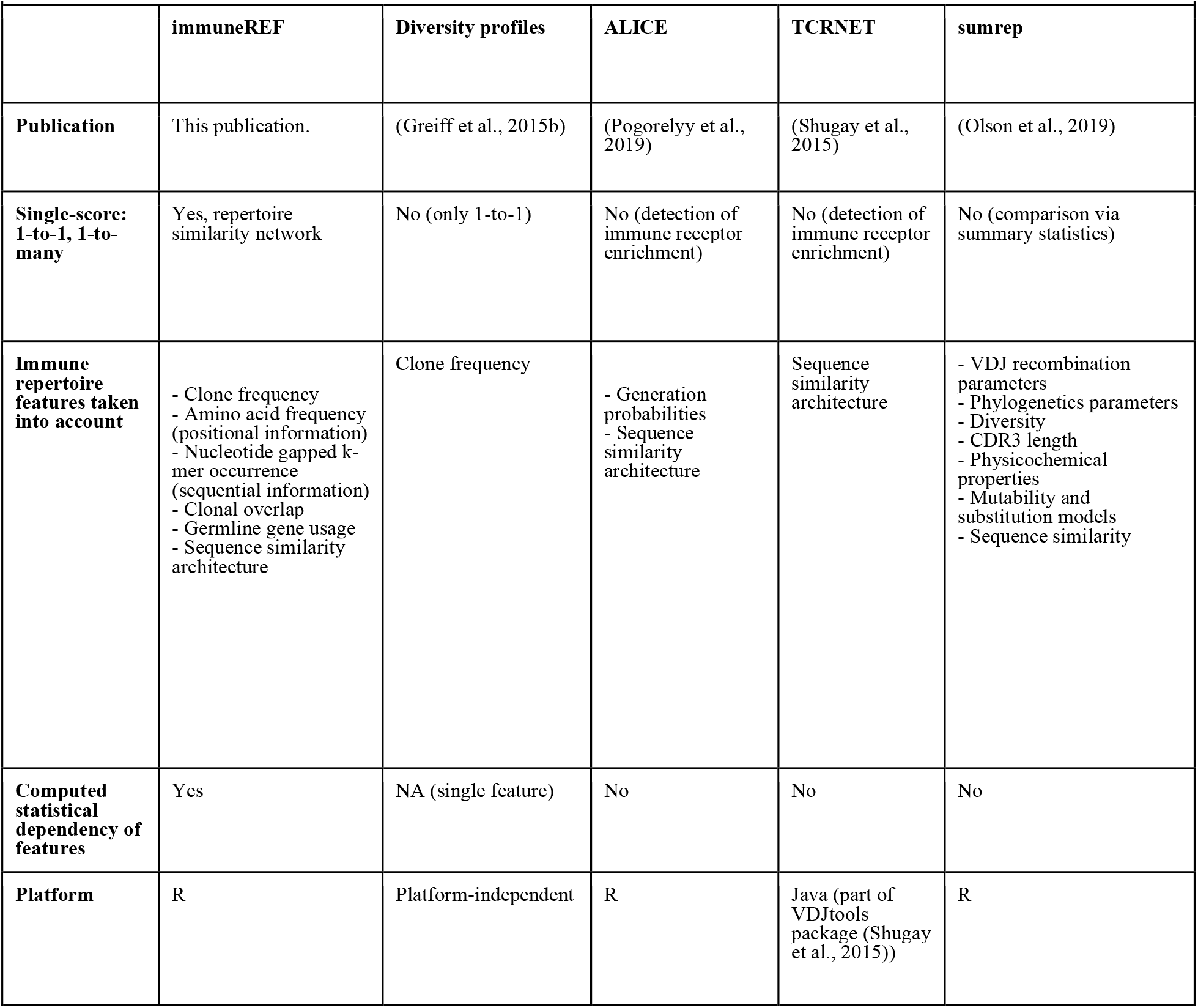
Comparison of immuneREF and other comparative methods. This table provides an overview of immuneREF features compared to previous comparative methods. Notably, immuneREF is the first instance of a comparative framework that takes into account multiple features of moderate to low statistical dependency to calculate a single repertoire score.

| Supplementary Table 4. Comparison of immuneREF and other methods for AIRR comparison |  |  |  |  |  |
| --- | --- | --- | --- | --- | --- |
|  | immuneREF | Diversity profiles | ALICE | TCRNET | sumrep |
| **Publication** | This publication. | (Greiff et al., 2015b) | (Pogorelyy et al., 2019) | (Shugay et al., 2015) | (Olson et al., 2019) |
| **Single-score: 1-to-1, 1-to-many** | Yes, repertoire similarity network | No (only 1-to-1) | No (detection of immune receptor enrichment) | No (detection of immune receptor enrichment) | No (comparison via summary statistics) |
| **Immune repertoire features taken into account** | - Clone frequency<br>- Amino acid frequency (positional information)<br>- Nucleotide gapped k-mer occurrence (sequential information)<br>- Clonal overlap<br>- Germline gene usage<br>- Sequence similarity architecture | Clone frequency | - Generation probabilities<br>- Sequence similarity architecture | Sequence similarity architecture | - VDJ recombination parameters<br>- Phylogenetics parameters<br>- Diversity<br>- CDR3 length<br>- Physicochemical properties<br>- Mutability and substitution models<br>- Sequence similarity |
| **Computed statistical dependency of features** | Yes | NA (single feature) | No | No | No |
| **Platform** | R | Platform-independent | R | Java (part of VDJtools package (Shugay et al., 2015)) | R |

**Supplementary Figure 1.**
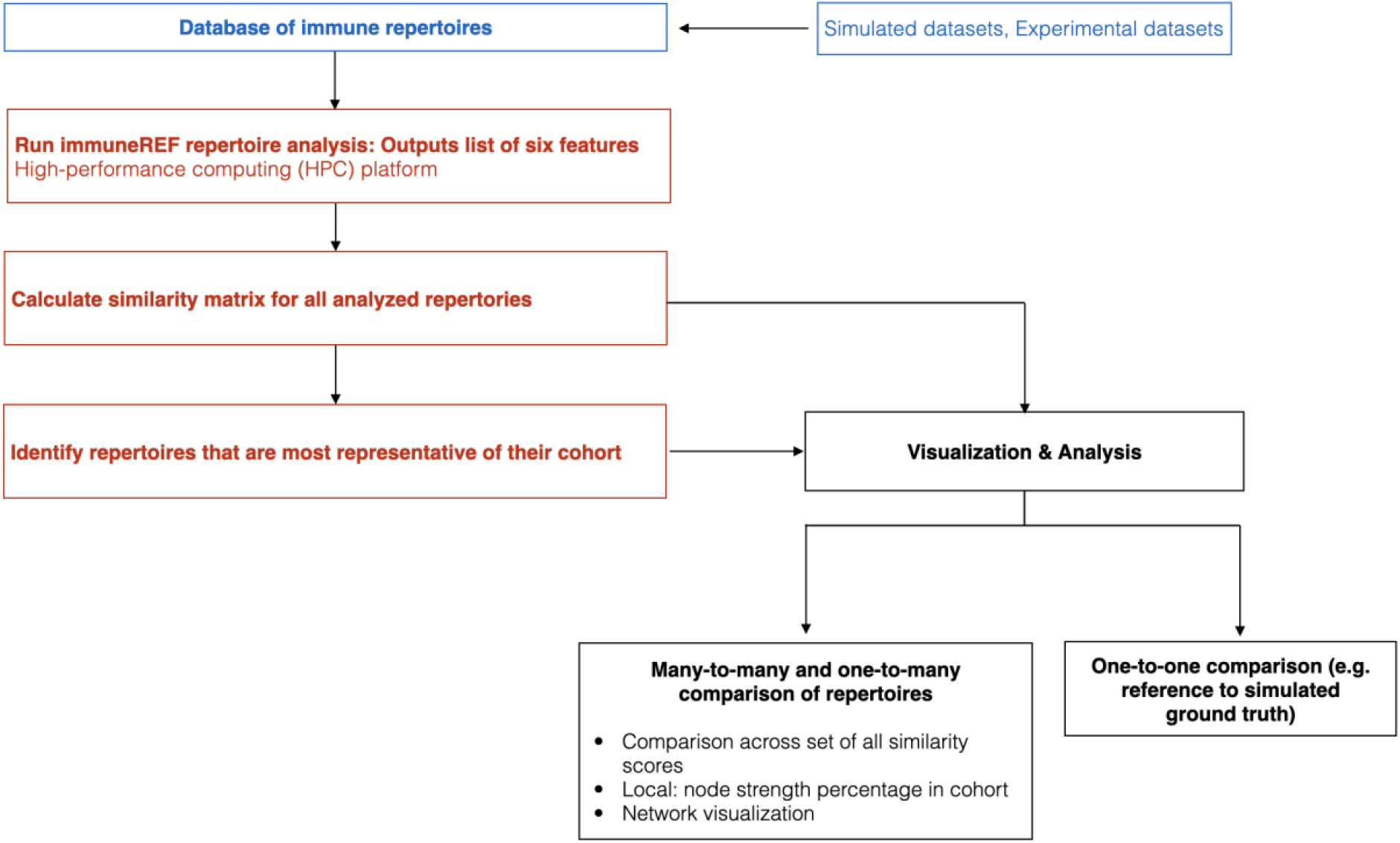
Flowchart of immune receptor repertoire analysis in the R package immuneREF. A database of repertoires (BCR/TCR, human/murine, simulated/experimental) is analyzed with respect to the six immuneREF features and their similarity scores are calculated. Repertoires that show the highest local similarity may be identified as representative for their cohort. The resulting similarity network may be analyzed in a many-to-many, one-to-many and one-to-one comparison.

**Supplementary Figure 2.**
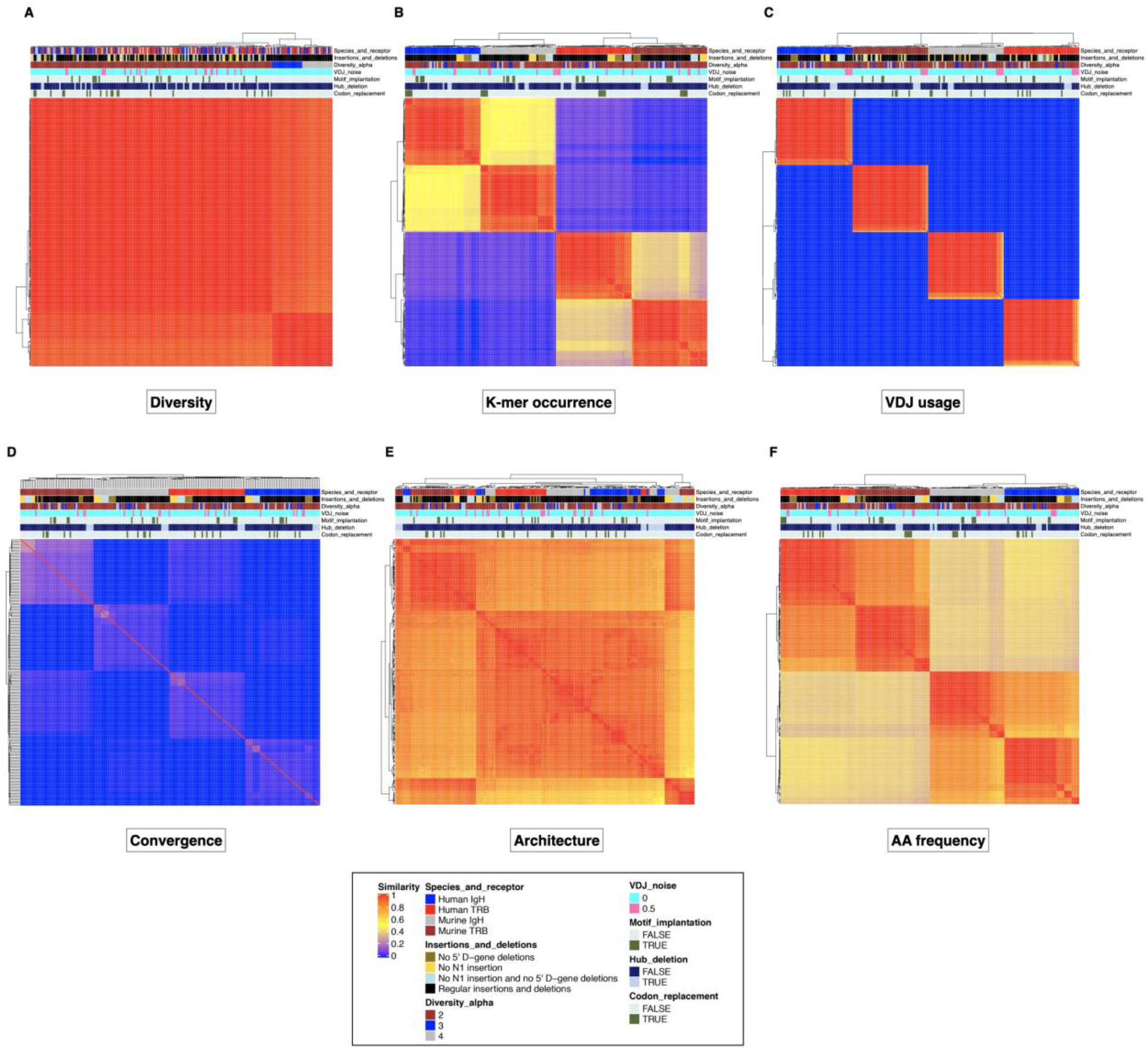
Every single feature of the similarity network shows a unique topology. (A–F) Heatmaps represent the similarity score landscape for each of the six features.

**Supplementary Figure 3.**
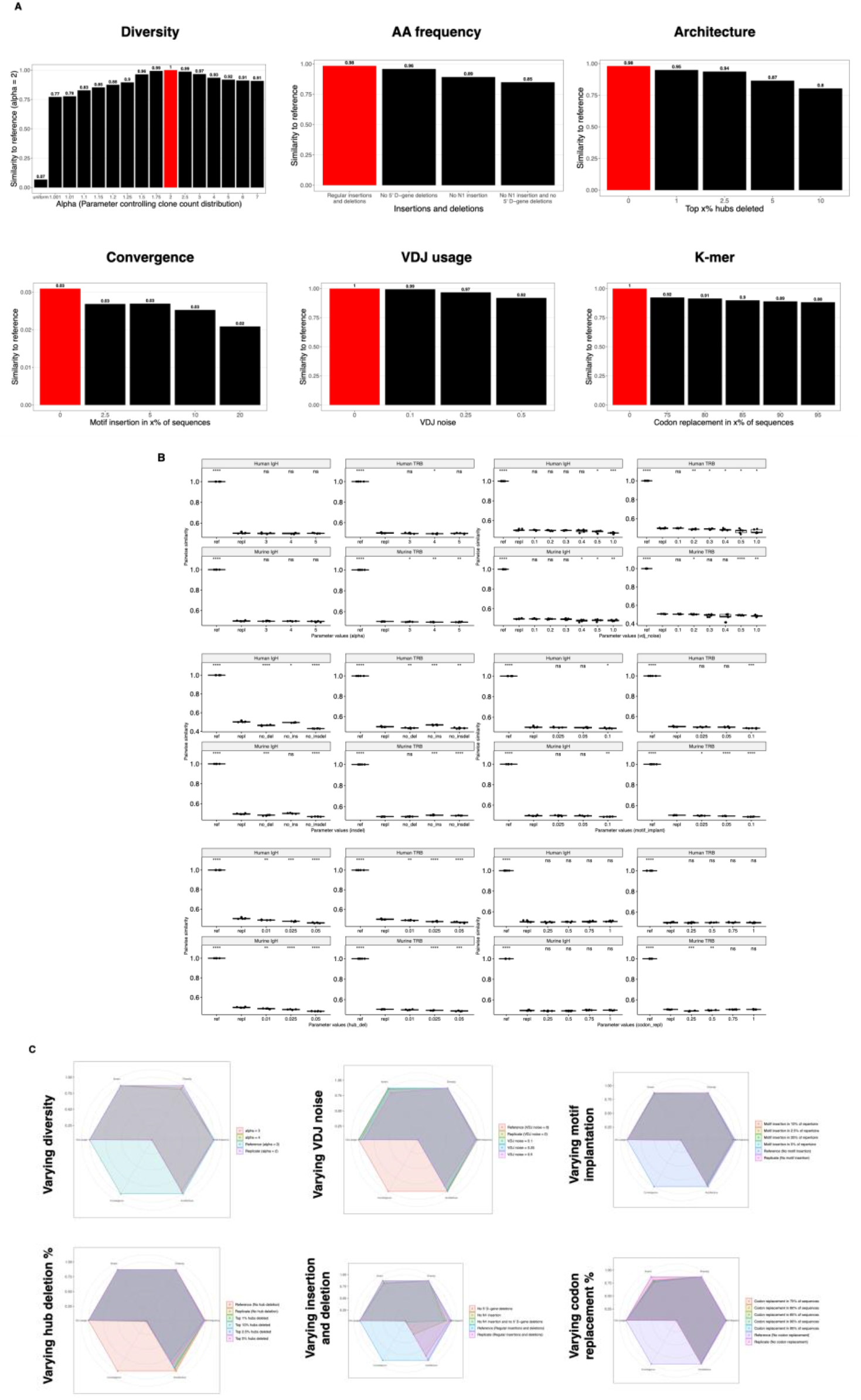
The similarity landscape of simulated repertoires defines a reference map for immune repertoire similarity. (A) Per feature differences for a range of immuneSIM parameters (red bar indicates default parameters) (see also SuppFig 3B,C) (B) Similarities in each feature across different parameters (see also Fig 3A) (C) Sensitivity of condensed similarity scores (y-axis) in response to different parameters (x-axis) faceted by species/receptor combination.

**Supplementary Figure 4.**
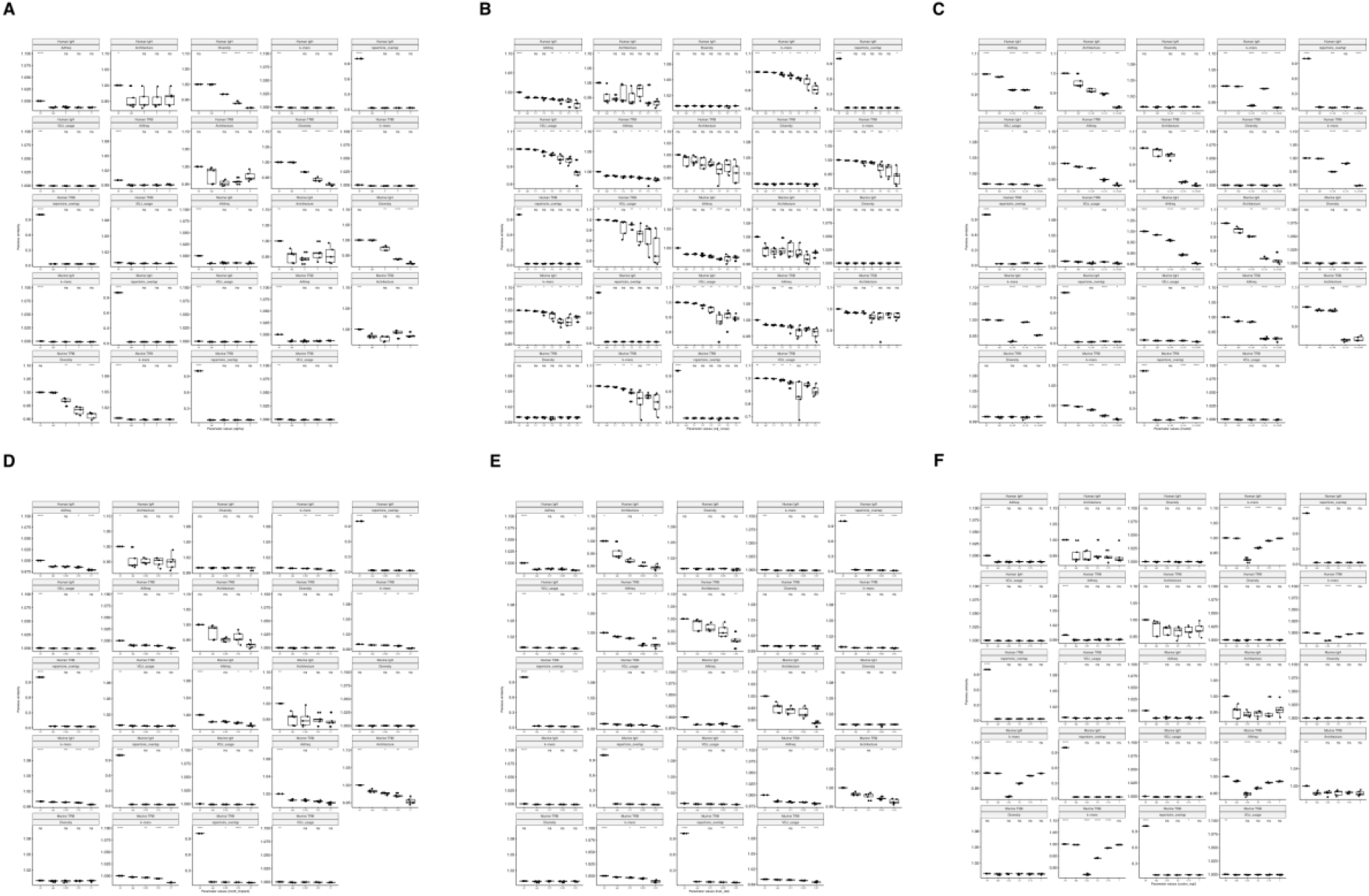
Sensitivity analysis per feature demonstrates the impact of each parameter change is feature specific. (A–F) The impact on pairwise similarity scores (y-axis) of parameter changes (x-axis) is shown for each parameter. Faceted by species/receptor combination and feature.

**Supplementary Figure 5.**
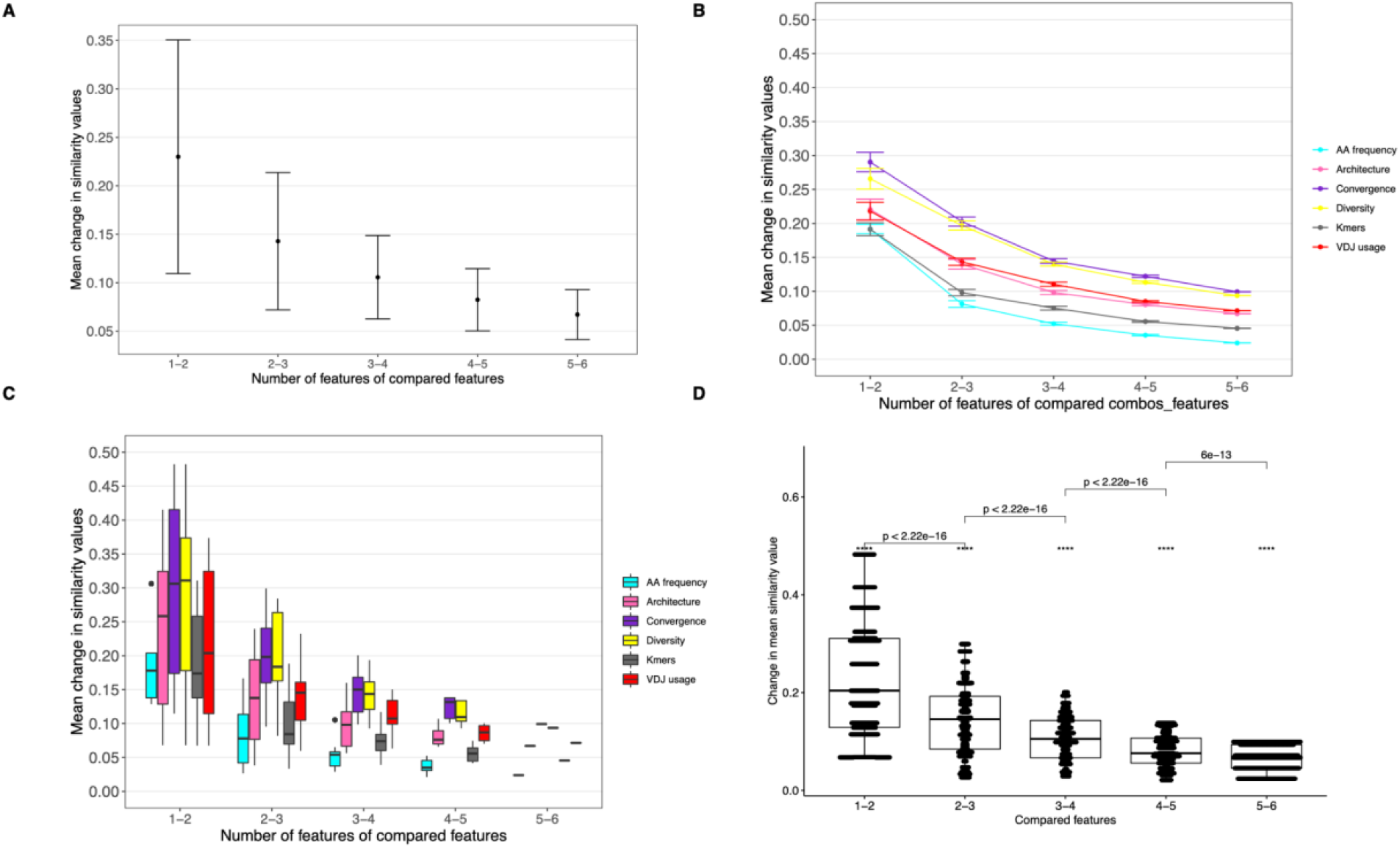
Diminishing marginal utility of additional features. (A) The mean change in similarity values after the addition of the next similarity feature (feature to be added is chosen randomly, 500 iterations) (B) The mean change in similarity values for the addition of each feature during various stages of the construction of the multi-feature network. (C) Boxplots of mean change in similarity values for the addition of each feature during various stages of constructing a multi-feature network. (D) Beeswarm representation of Fig. 2D.

**Supplementary Figure 6.**
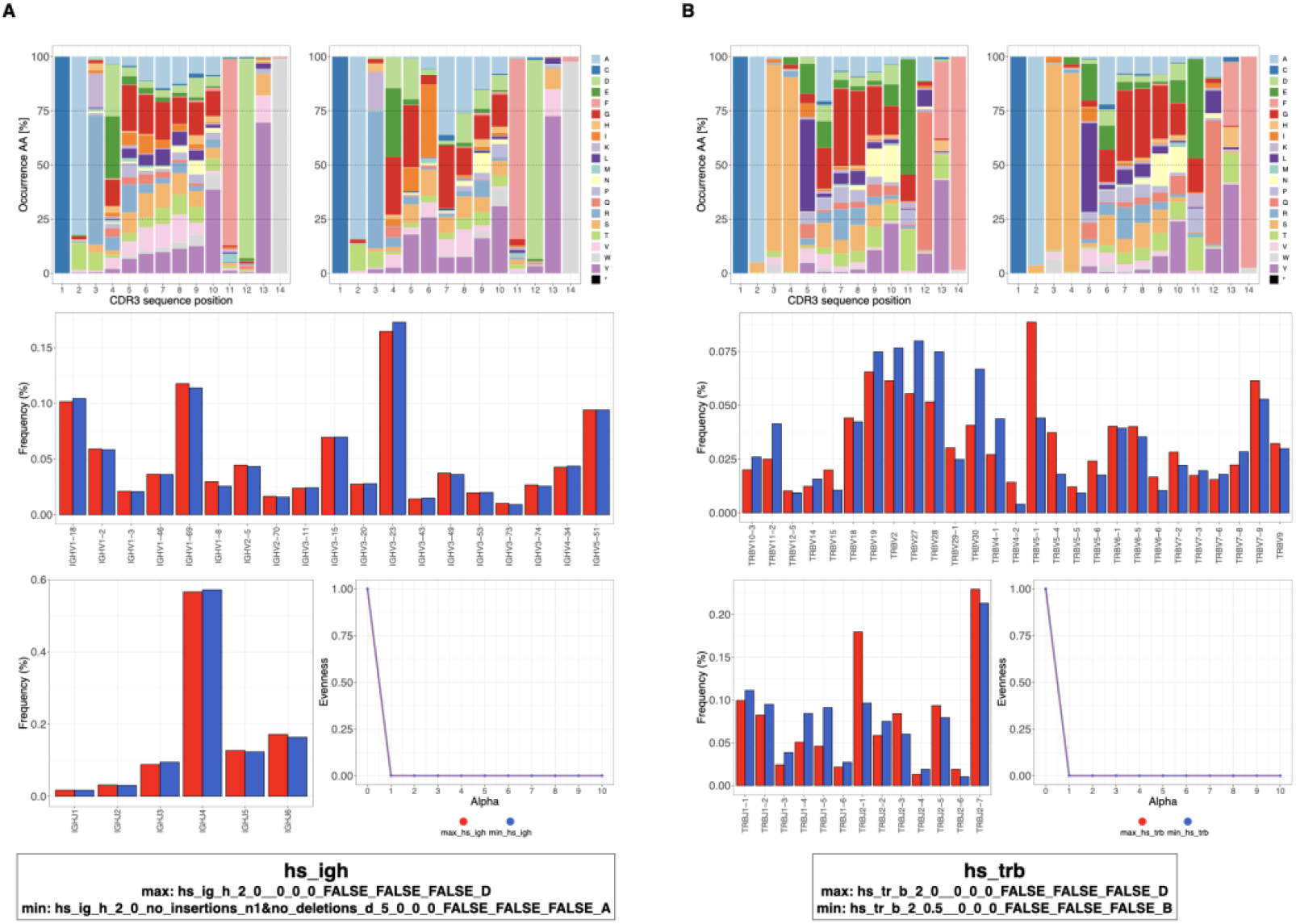
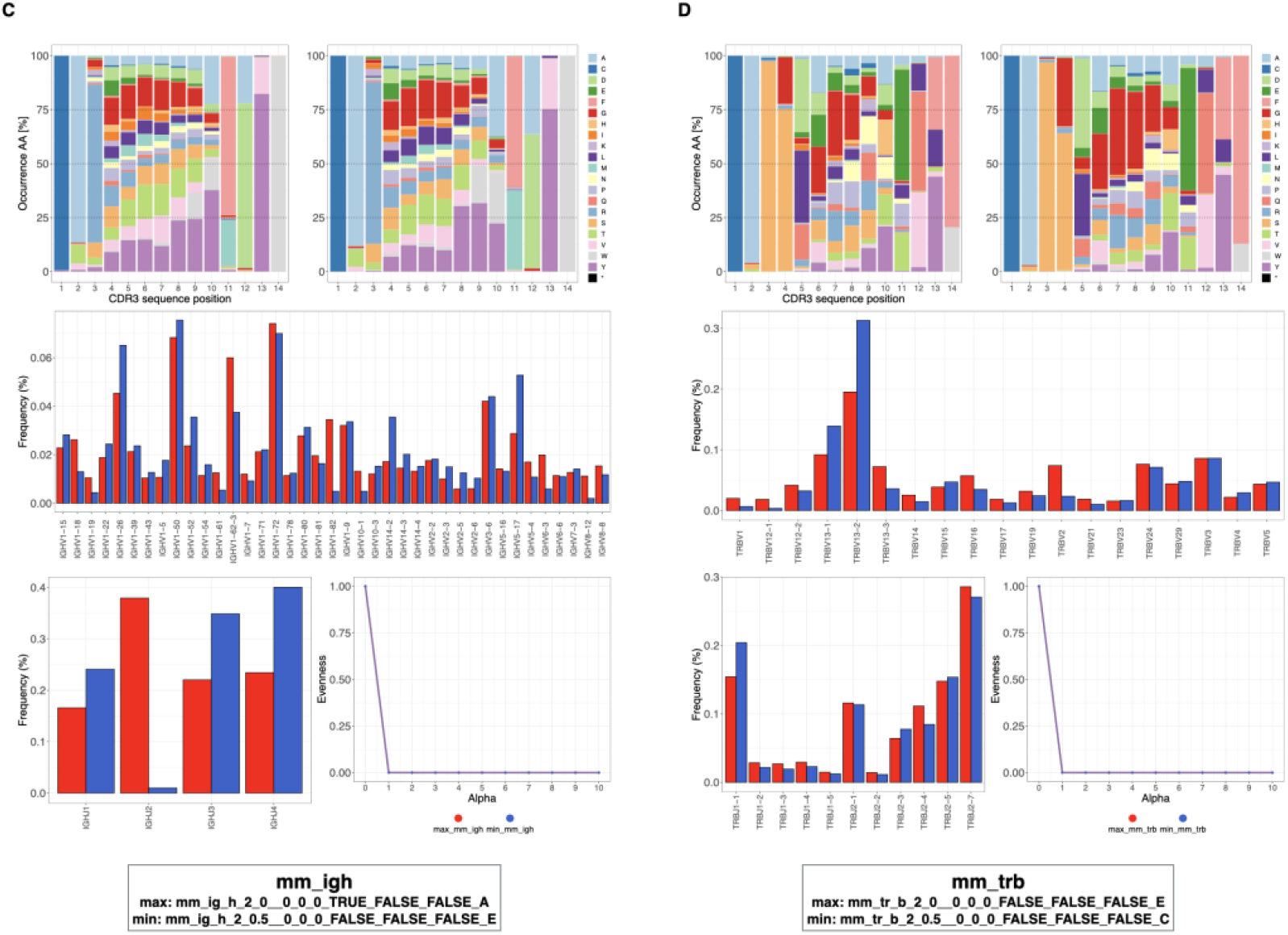
Within-cohort variability of simulated repertoires varies by receptor cohort (human and murine simulated repertoires). The amino acid frequency (first row), VJ usage (middle row, bottom left), and evenness (bottom right) are compared between the most and least locally similar repertoires of human (hs) **(A)** IgH and **(B)** TRB simulated repertoires as well as mouse (mm) **(C)** IgH and **(D)** TRB simulated repertoires.

**Supplementary Figure 7.**
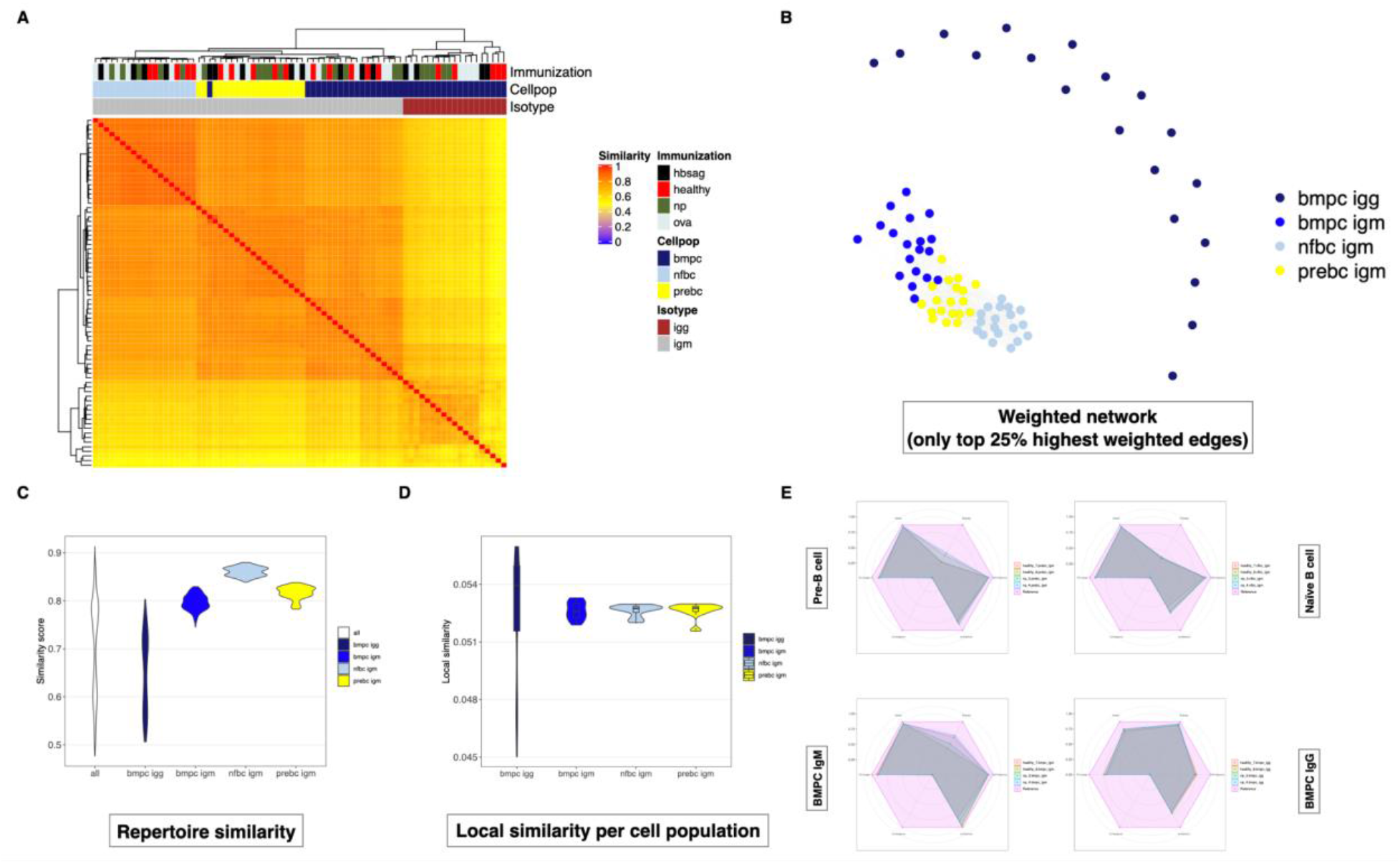
Application of immuneREF to 76 experimental repertoires from mouse immunization study (Greiff et al. 2017a) finds differences in cell population/isotype cohorts. (A) Similarity landscape of experimental (murine, BCR) repertoires from four immunization cohorts (Healthy, HBsAg, NP, OVA) and four cell populations (pre-B cell IgM, naïve B-cell IgM, bone marrow plasma cell IgM and IgG) (B) Network visualization of the 76 nodes and weighted edges between repertoires of similarity scores (top 25% edge by edge weights). (C) Distribution of similarity scores across the entire network and per cell population shows variation within and across cell populations. (D) Distribution of local similarity values per repertoire for each cell population. (E) Comparison of most locally similar repertoires of the healthy and NP cohort for each cell population and an immuneSIM reference repertoire (standard immuneSIM parameters).

**Supplementary Figure 8.**
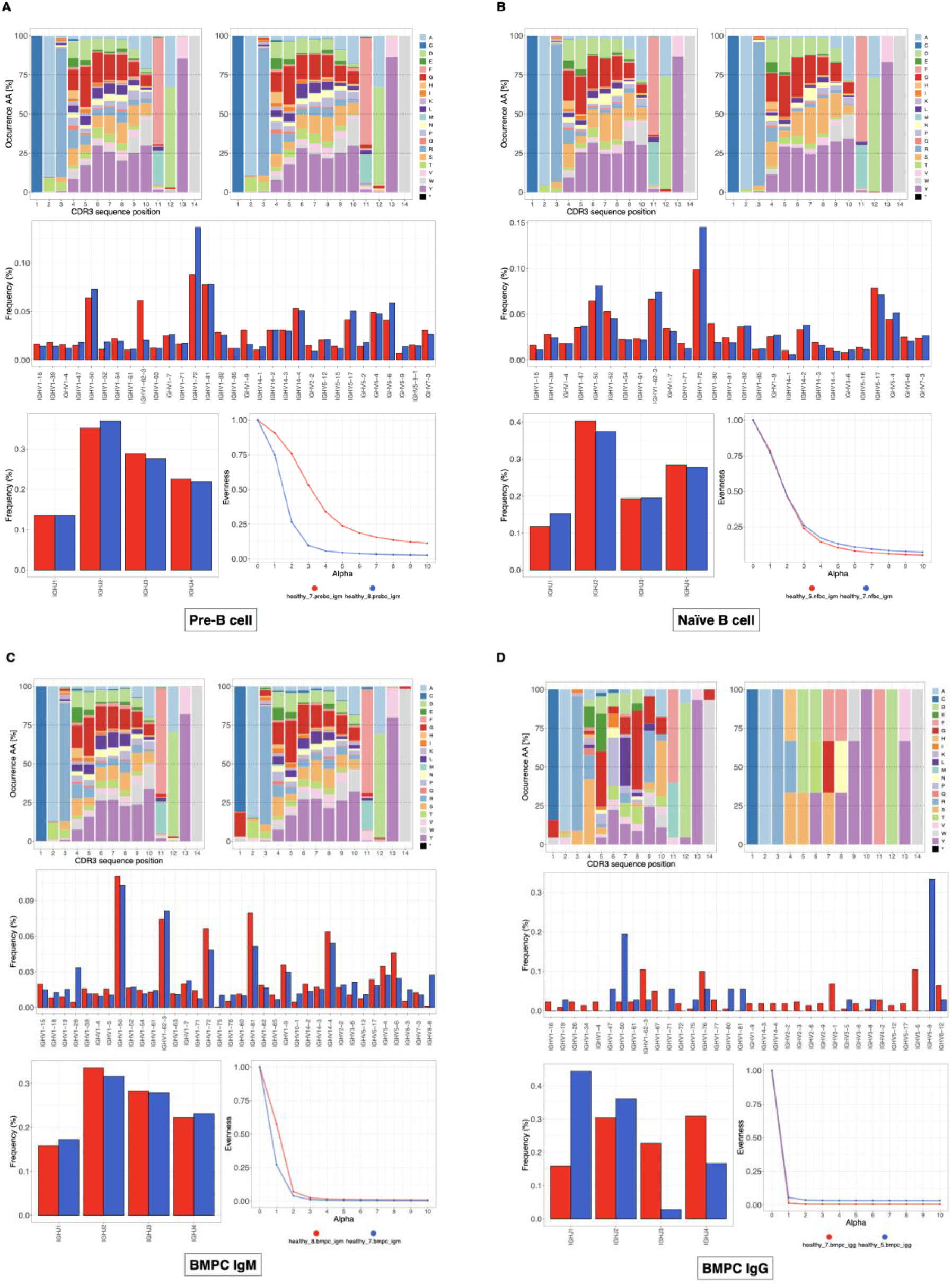
IgM isotype repertoires of the mouse B-cell dataset (Greiff et al., 2017a) have high within-cell population similarity. The amino acid frequency (first row), VJ usage (middle row, bottom left), and evenness (bottom right) are compared between the most and least locally similar repertoires of each cell population in the healthy cohort.

**Supplementary Figure 9.**
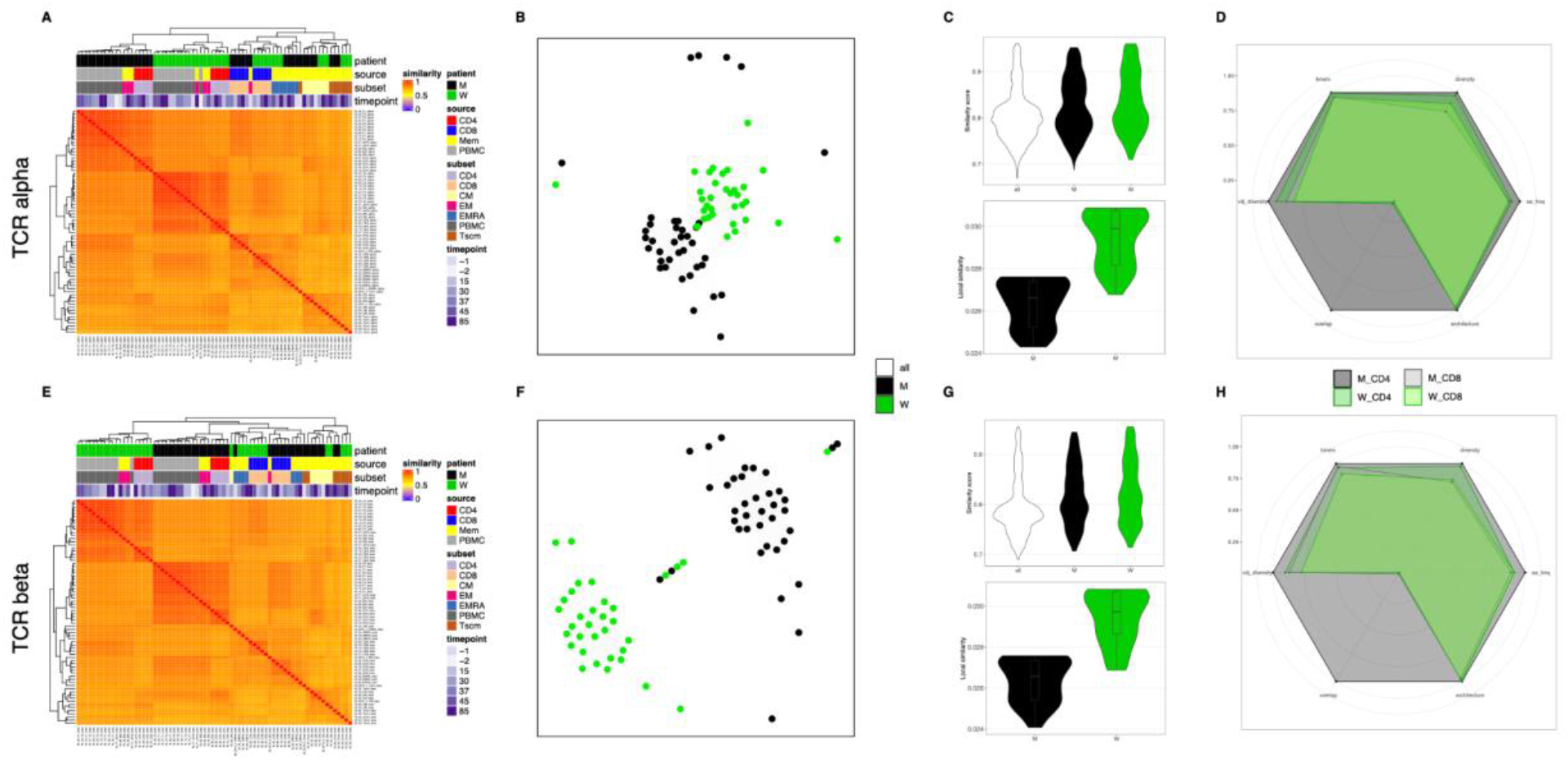
Application of immuneREF to 72 T-cell repertoires from various cell populations of two Covid patients (longitudinal). (A,E) Similarity landscape of experimental (human, TCR, alpha and beta) repertoires for four cell populations (B,F) Network visualization of the 72 nodes and weighted edges between repertoires of similarity scores (top 25% edge by edge weights). (C,G) Distribution of similarity scores across the entire network (top) and local similarity scores (bottom) per patient. (D,H) Comparison of most and locally similar repertoires of each patient from the CD4 and CD8 subsets.

**Supplementary Figure 10.**
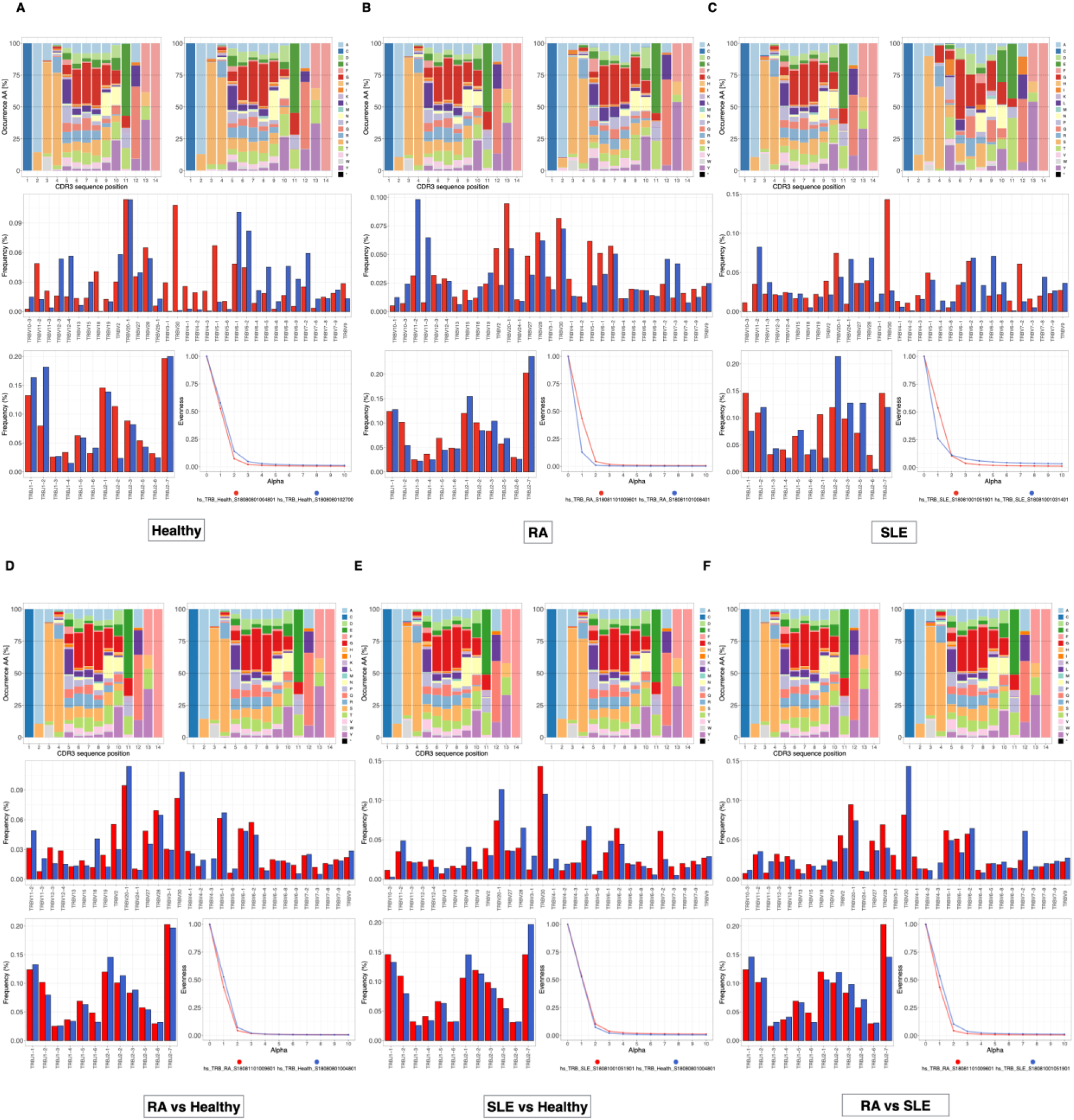
Within and cross-immune state similarity of repertoires is high (Healthy, RA, SLE) for the PIRD human T-cell dataset (Zhang et al.,2019). The amino acid frequency (first row), VJ usage (middle row, bottom left) and evenness (bottom right) compared (A–C) between the most (left amino acid plot, red bars, and lines) and least (right amino acid plot, blue bars, and lines) locally similar repertoires of each immune state. (D–F) between the most locally similar repertoires of each immune state pair.

**Supplementary Figure 11.**
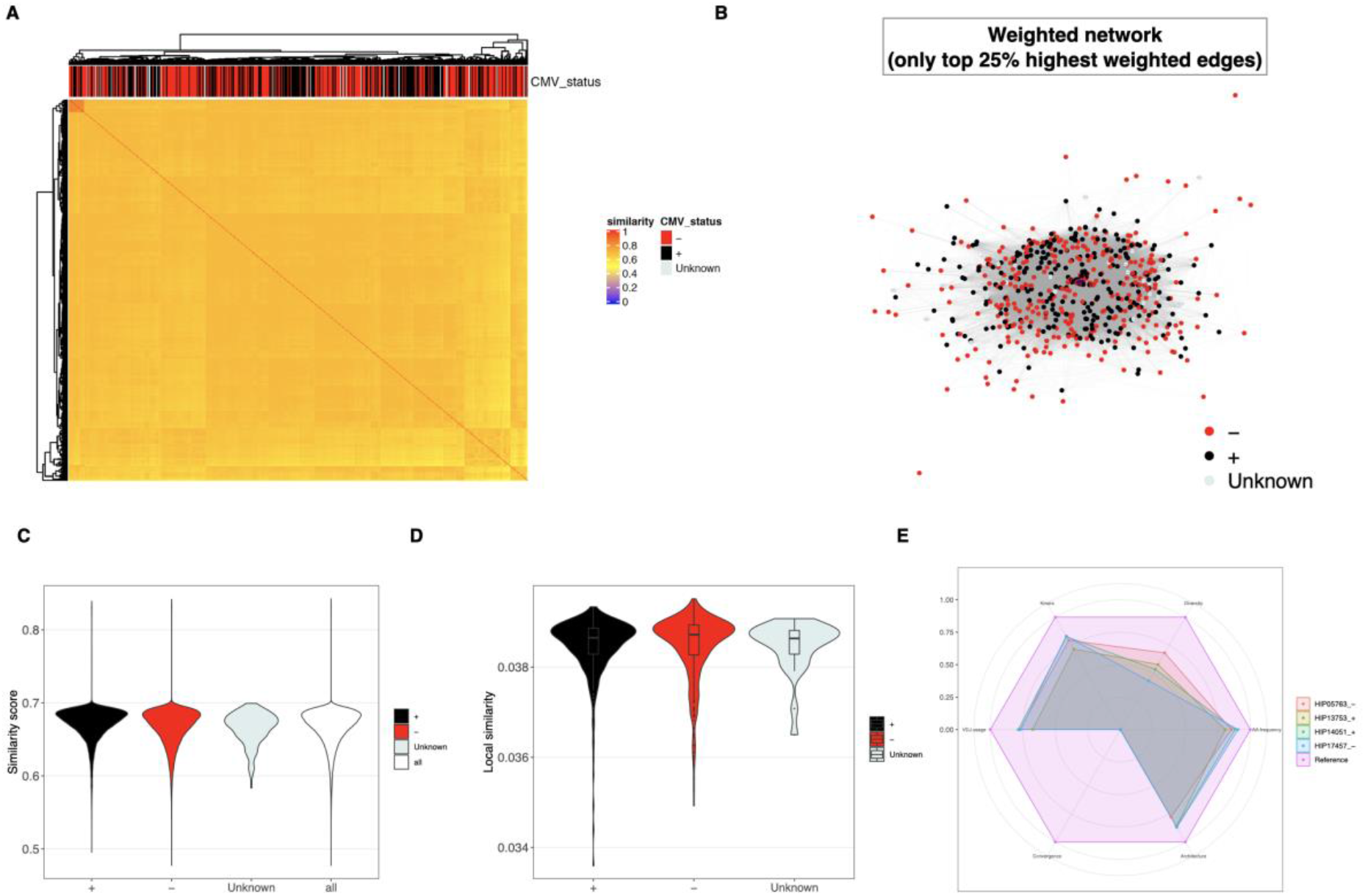
Application of immuneREF to 666 experimental, CMV-serotyped repertoires (human, TCR, Emerson et al., 2017) finds even similarity distribution across repertoires of CMV+ and CMV- status. (A) Similarity landscape of experimental (human, TCR) repertoires of CMV+ (289), CMV- (352) and unknown serotype (25) (B) Network visualization of the 666 nodes and weighted edges between repertoires of similarity scores (top 25% edge by edge weights). (C) Distribution of similarity score across the entire network and per immune state shows similar distribution CMV+ and CMV- population. (D) Distribution of local similarity values per repertoire for each serotype. (E) Comparison of most and least locally similar repertoires of the CMV+ and CMV- cohort.

**Supplementary Figure 12.**
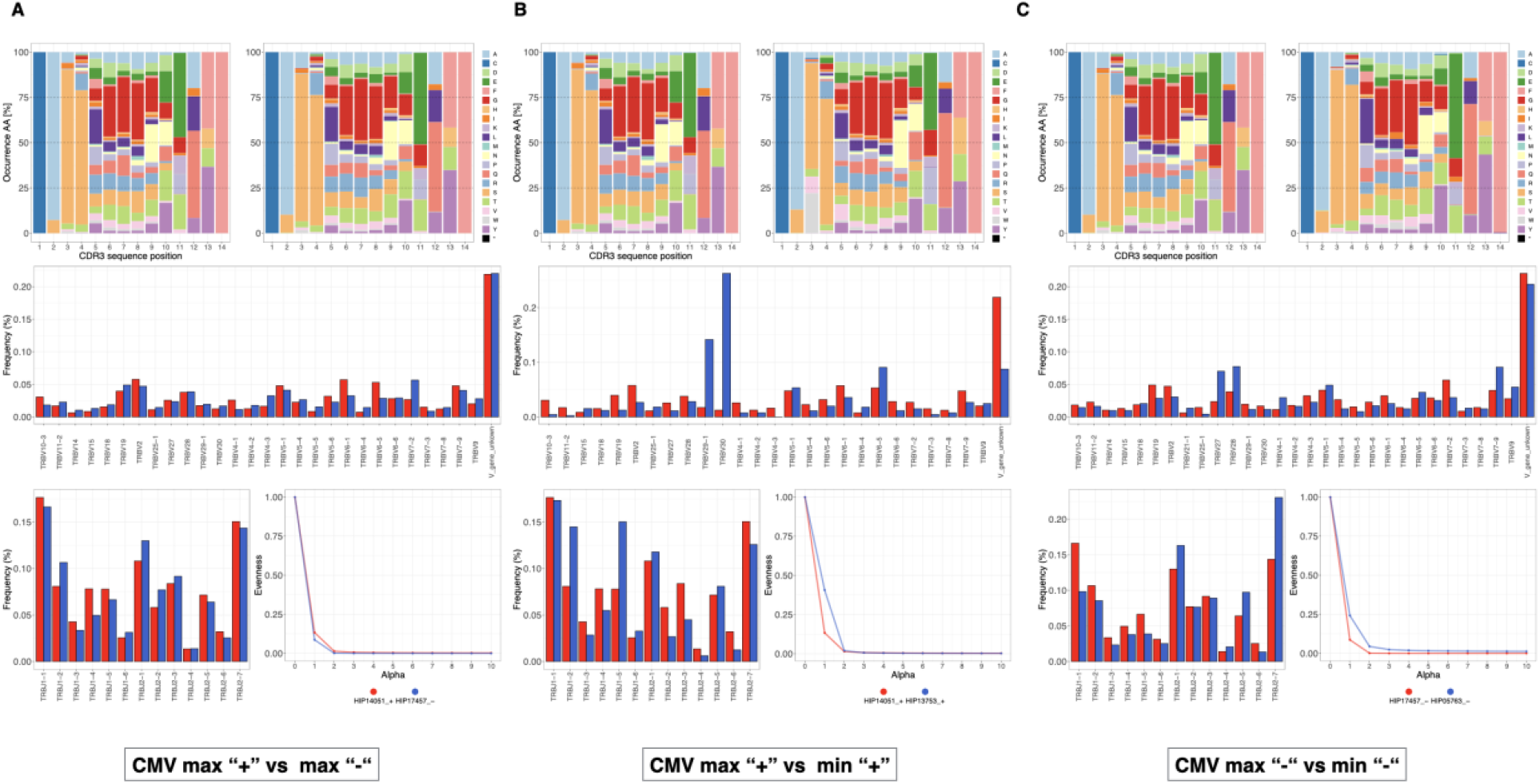
Within and across immune state similarity of repertoires is high for the CMV dataset (human, TCR, Emerson et al. 2017). The amino acid frequency (first row), VJ usage (middle row, bottom left), and evenness (bottom right), compared between (A) the most and locally similar repertoires of CMV+ and CMV- cohorts. (B) CMV+ and (C) CMV- repertoires with the lowest and highest local similarity.

**Supplementary Figure 13.**
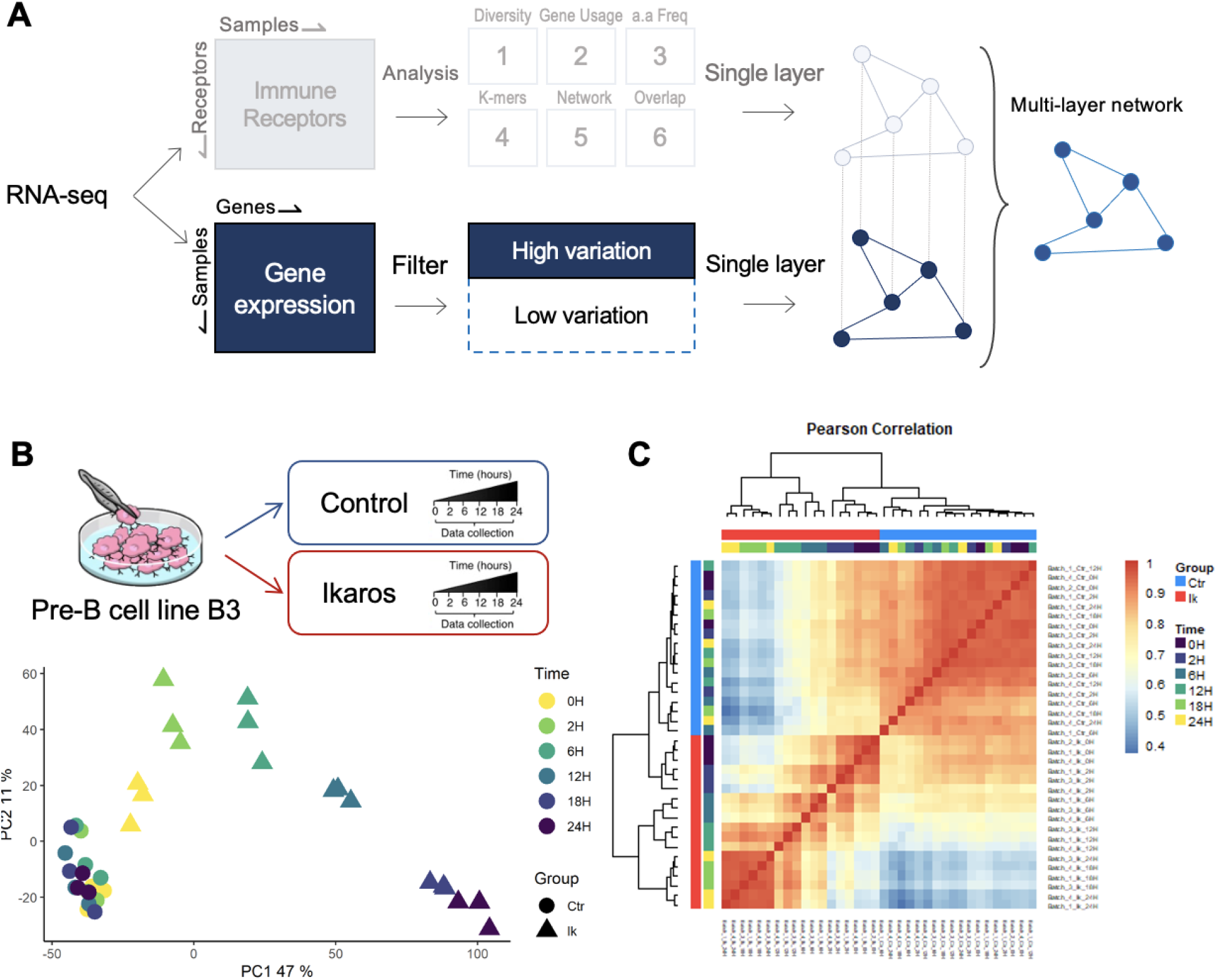
Gene expression and immune repertoire integration. Gene expression can be added as an additional single feature to obtain the multi-feature network. (A) Pipeline overview. (B) Example of analysis using STATegra dataset (mouse pre-B cell line B3, RNA-seq, Gomez-Cabrero et al. 2019). (C) Heatmap representing co-expression patterns between samples.

